# Time, Space and Single-Cell Resolved Molecular Trajectory of Cell Populations and the Laterality of the Body Plan at Gastrulation

**DOI:** 10.1101/2023.04.19.537134

**Authors:** Ran Wang, Xianfa Yang, Jiehui Chen, Lin Zhang, Jonathan A. Griffiths, Guizhong Cui, Yingying Chen, Yun Qian, Guangdun Peng, Jinsong Li, Liantang Wang, John C. Marioni, Patrick P.L. Tam, Naihe Jing

**Author notes:** Correspondence (P.P.L.T.), (N.J.).

## Abstract

Understanding of the molecular drivers of lineage diversification and tissue patterning during primary germ layer development requires in-depth knowledge of the dynamic molecular trajectories of cell lineages across a series of developmental stages of gastrulation^1–7^. Through computational modeling, we constructed at single-cell resolution a spatio-temporal compendium of the molecular trajectories of germ-layer derivatives in gastrula-stage mouse embryos. This molecular atlas infers the developmental trajectories of single-cell populations and the molecular network activity underpinning the specification and differentiation of the germ-layer lineages. Analysis of the heterogeneity of cellular composition of cell populations at defined positions in the epiblast revealed progressive diversification of cell types, mirroring the process of lineage allocation during gastrulation. A novel observation is the difference in the contribution of cells on contralateral sides of the epiblast to mesoderm derivatives of the early organogenesis embryo, and the enhanced BMP signaling activity in right-side mesoderm of E7.5 embryo. Perturbation of BMP signaling activity at late gastrulation led to randomization of left-right (L-R) molecular asymmetry in the lateral mesoderm of early-somite-stage embryo. Our findings indicate the asymmetric BMP activity during gastrulation may be critical for the symmetry breaking process associated with specification of L-R body asymmetry ahead of the acquisition of functionality of the L-R organizer.

---

The molecular architecture of cell populations in mouse embryos at pre-to late-gastrulation stages has been charted by profiling the transcriptome of small groups of cells captured from defined locations in the germ layers^1^. These data have provided insight into the developmental trajectory of the germ layer tissues and the activity of the molecular networks associated with the transition of pluripotency states, and the specification and regionalization of the germ layer derivative during gastrulation. Since the transcriptomic profile was collated from populations of cells, it is not feasible to resolve the divergence in transcriptome activity between cells of different identities within each population. Recently, single-cell genomics and transcriptomics have been applied to investigate the molecular attributes and potential lineage relationship of the multitude of cell types in the embryo during development^2, 8^. However, profiling the population of single cells pooled from embryos, where spatial information is lost, and analyzing cells from embryos at one developmental time point cannot provide insight into the molecular architecture of individual cells that are regionalized to specific compartment (space) in the embryo across the developmental stages (time). It is therefore imperative to combine existing spatially resolved transcriptome information with single-cell transcriptomes to construct a high-dimension, in time and space (4D), molecular atlas of spatially-and stage-resolved single-cell gene expression profiles of cells in the germ layers of the embryo. The 4D molecular atlas so constructed will enable prediction of the developmental trajectories of germ layer tissues and the inference of molecular drivers underpinning the diversification of tissue lineages and the molecular activity associated with lineage development and tissue patterning during gastrulation.

## The spatio-temporal molecular atlas

A key prerequisite for the construction of the atlas of molecular trajectories of cells in the embryo is the collation of a stage-and spatially-registered population-based transcriptome dataset. To this end, we have enriched the transcriptome data of mouse embryonic day (E) 6.5-7.5 embryos, by capturing two additional time points: E6.75 and E7.25, and integrating the data into the established spatio-temporal transcriptome^1^. The full transcriptome dataset comprises of Geo-seq data of 29 (E6.5), 29 (E6.75), 73 (E7.0), 94 (E7.25) and 81 (E7.5) positionally registered samples from the epiblast, ectoderm, mesoderm and endoderm of embryos (in replicates, Extended Data Fig. 1a-c). The data were rendered digitally for depiction in 2D corn plots, with the samples staged by the developmental status of the primitive streak (by *T* expression) and the ectoderm progenitor (*Otx2* expression) (Extended Data Fig. 1d). Quality assessment of this dataset showed that a median of about 11,000 genes were detected per sample across all embryos (Extended Data Fig. 1e, f), with an average of 10 million reads per library that ensured the sufficient sequencing depth saturation (Extended Data Fig. 1g).

**Figure 1.**
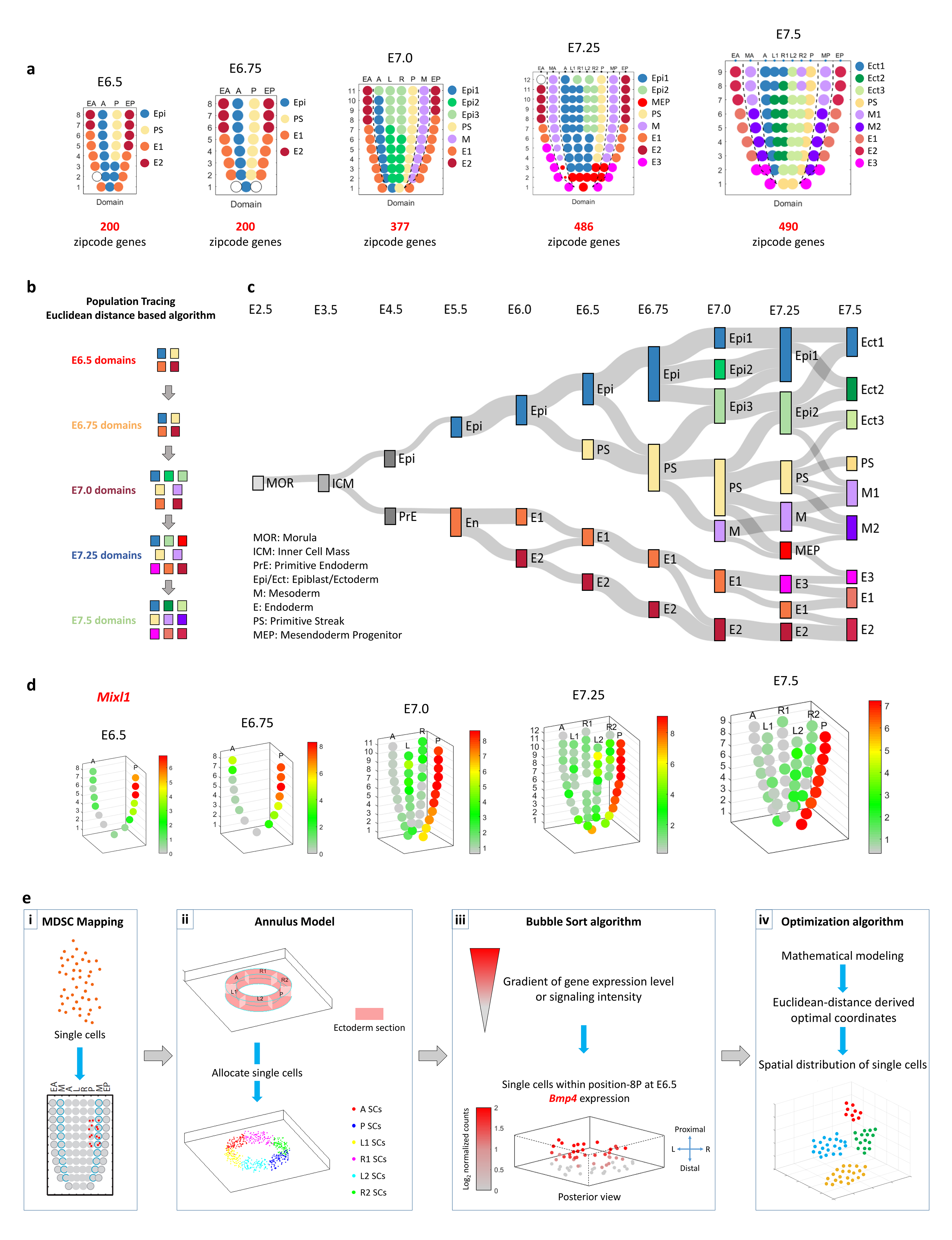
Generation of a spatio-temporal molecular atlas of the germ layers of gastrula-stage mouse embryo. **a.** Spatial domain of cell populations in the epiblast/ectoderm, mesoderm and endoderm of E6.5-E7.5 embryos, defined by the position-specific expression of zipcode gene transcripts. Geo-seq sampling positions: epiblast/ectoderm – A, anterior; L, left lateral; R, right lateral; L1/R1, left/right anterior lateral, L2/R2, left/right posterior lateral; M, mesoderm – MA, anterior mesoderm; MP, posterior mesoderm; E, endoderm – EA, anterior endoderm; EP, posterior endoderm. Number: descending series indicating positions in the proximal-distal axis. Germ layer domains: Epi: epiblast, Epi1, 2, 3: epiblast domain 1, 2 and 3; M: mesoderm, M1, M2: mesoderm domain 1 and 2; MEP, putative mesendoderm progenitors; E: endoderm, E1, E2, E3: endoderm domain 1, 2 and 3; PS, primitive streak. **b.** The structure of the Population Tracing algorithm for imputing the developmental connectivity of cell populations across stages of gastrulation (see detail of mathematical operations in Methods and Extended Data Fig. 2f). **c.** The developmental trajectory of sub-populations within each germ layer tissue domains descending from blastomeres of the preimplantation E2.5 morula stage embryo to the germ layers of E7.5 late-gastrulation stage embryo. **d.** 3D Model of the epiblast/ectoderm displaying the cell populations by imputed positional coordinates (see Methods for detail of the mathematical modeling). The exemplar 3D corn plots show the spatio-temporal distribution of *Mixl1*-expressing population in the primitive streak, the proximal-distal span of the *Mixl1*+ domain defines the developmental stage of the gastrulating embryo. The color legend indicates the level of expression determined by the transcript counts. **e.** A flow diagram of the 4-step spatial mapping protocol. 1) Multi-Dimension Single-Cell (MDSC) Mapping allocates single cells to their imputed position. 2) Annulus Model simulates the Geo-seq positions. Single cells that mapped to a Geo-seq position were distributed uniformly across the interior space of the position in each annulus section. 3) Bubble Sort algorithm displays the cells in relation to the gradient of gene expression level or signaling intensity. 4) Optimization algorithm refines and visualizes the spatial distribution pattern of cell types by optimal coordinates at each Geo-seq position.

The spatio-temporal transcriptome can be mined to identify the position-specific signature genes of populations of cells in different domains of the germ layers of the embryos. Applying BIC-SKmeans and PC-loading analysis, the number of distinct gene-expression domains (Extended Data Fig. 2a-e) and the position-specific signature genes for corn plot (i.e., Geo-seq) position (Fig. 1a; designated as the zipcode genes, zipcodes for short)^9–11^ were identified in the germ layers of E6.5-E7.5 embryos (see Methods). To trace the developmental connectivity of cell populations in various domains, we devised a Population Tracing algorithm (Fig. 1b, Extended Data Fig. 2f and Methods) for inferring the putative molecular trajectory of cell populations at successive developmental stages. The algorithm was applied by collating the zipcodes of cell population at different developmental stages as the input gene set, and calculate the Euclidean distance of any two Geo-seq domains in embryos of successive developmental stages, to identify the most closely-connected domains. Using a combined dataset of the transcriptome of pre-and peri-implantation embryo^12^ and the E5.5-E7.5 embryo^1^, we charted the developmental trajectory of cell populations from blastomeres of the morula to the nine major germ layer tissue domains at E7.5 (Fig. 1c). The developmental trajectory has also provided insights into the location of unique progenitor cell types and the molecular activity that may be related to their ontogeny and differentiation. For example, through PC loading analysis, a mixed cell population containing putative mesendoderm progenitor (MEP) (Fig. 1a) was pinpointed by the expression of Group (G)-5 and G-7 genes to the E7.25 anterior primitive streak (PS) and the adjacent mesoderm (Extended Data Fig. 2d). By E7.5, the mixed population of dual potential precursor cells diverged into two cell types, the distal mesoderm (M2) and distal endoderm (E3) (Fig. 1c). Another example arose from tracking the expression of signature genes is the emergence of the node that is marked by *Noto* and *Foxj1* (Extended Data Fig. 2d, g), and the functional ontogeny of gene groups (Extended Data Fig. 2h, i) that are characteristics of the gastrula organizer^13^. We also identified two cell populations in the proximal-lateral ectoderm (at positions: 8R2 and 9R2) of the E7.5 embryo (Extended Data Fig. 2e) that showed significant connectivity of transcriptome features (gene group G-8) with the mesoderm in the proximal region (M1, position 7- 9MA/P) (see details later). Moreover, the definition of spatial domains has a high inter-embryo consistency (Extended Data Fig. 3a-e), implicating a significant synchronicity in the patterning of cell populations in the germ layers during gastrulation.

**Figure 2.**
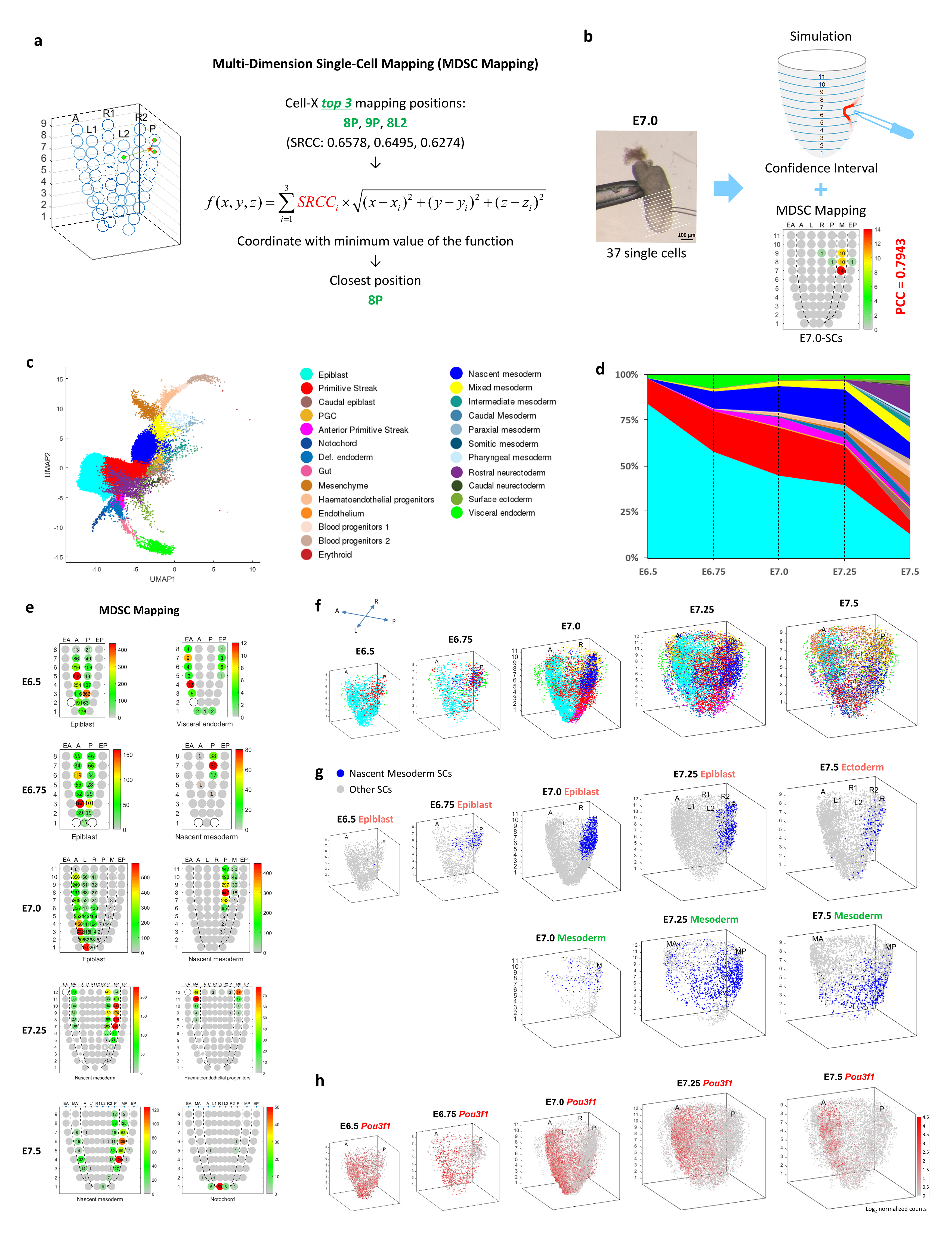
A single-cell resolution 4D molecular atlas of mouse gastrulation. **a.** The pipeline of Multi-Dimension Single-Cell (MDSC) Mapping. The SRCCs of the expression values of the zipcodes of each single cell against all reference samples of the reference embryo were calculated, followed by the application of a spatial smoothing algorithm to impute the high-confidence (closest) location (see Methods). The mapping of cells to position 8P was shown as an example. **b.** Verification of the results of MDSC Mapping of single cells isolated from a known position in E7.0 embryo. The number on each corn indicates the number of cells mapped to the Geo-seq position in the germ layers. PCC values and confidence intervals are shown for the simulation. **c.** Uniform manifold approximation and projection (UMAP) plot showing the data structure of the ‘Gastrulation Atlas’ comprising 32,940 cells from E6.5 to E7.5 embryos, with the exclusion of extraembryonic cells. Twenty-five cell types are annotated (see color legend). Def. endoderm, definitive endoderm; PGC, primordial germ cells. **d.** Fraction of cell type per time point, displaying a progressive increase in cell-type complexity during gastrulation. **e.** MDSC Mapping results of exemplar cell types for E6.5, E6.75, E7.0, E7.25 and E7.5 embryos. The number in each corn indicates the number of cells mapped to the specific Geo-seq position. **f.** The spatio-temporal distribution of all the single cells identified in the Gastrulation Atlas of E6.5-E7.5 mouse embryos. Cell types are annotated as in Fig. 2c. **g.** The spatio-temporal distribution of single cells annotated as “nascent mesoderm” in the epiblast and mesoderm of E6.5-E7.5 embryos. **h.** The spatio-temporal distribution of *Pou3f1*-expressing cells in E6.5-E7.5 embryos. The color legend indicates the level of expression determined by the transcript counts.

**Figure 3.**
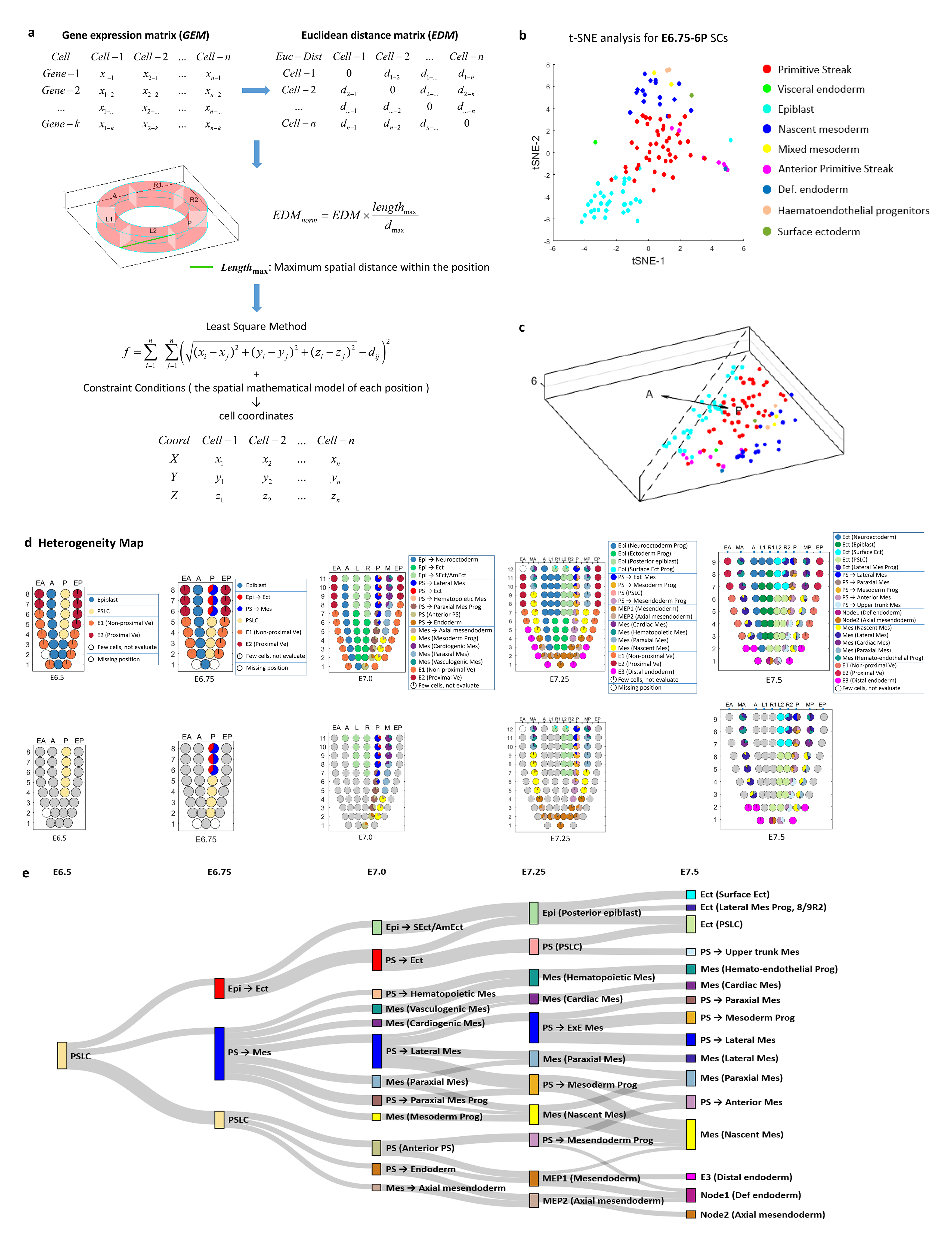
Mathematical modelling for single-cell spatial distribution and the collation of spatio-temporal heterogeneity map. **a.** The mathematical model for re-ordering the spatial distribution of the single cells within a Geo-seq position by Euclidean-distance derived optimal coordinates (see Methods). **b.** *t*-distributed stochastic neighbor embedding (*t*-SNE) plot showing the single cells mapped to position-6P at E6.75. Cell types are annotated (see legend). **c.** The imputed spatial distribution of single cell types within position-6P at E6.75. Cell types are annotated as in Fig. 3b. **d.** Heterogeneity Map. Corn plots (top row) show the composition of cell types (shown as pie chart in each corn) at different Geo-seq positions in the germ layers across the five timepoints of gastrulation. Corn plots (bottom row) show the heterogeneity of cell populations descending from the primitive steak like cells in different Geo-seq position in the germ layers of the gastrulation stage embryo. **e.** Molecular trajectory of the descendants of primitive streak like cells of the E6.5 posterior epiblast in the germ layers of E6.75, E7.0, E7.25 and E7.5 embryo, imputed using the Population Tracing algorithm. Cell types are listed in Extended Data Fig. 8. The rectangle represents the spatialized cell type (color indicating the cell types) with the size indicating the propensity of branching trajectory, and width of the edge indicates the strength of correlation between the connected cell types.

To mimic the spatial structure and visualize the spatial pattern of gene expression in the embryo, the spatio-temporal transcriptome data were then rendered digitally for depiction in ‘3D corn plots’ (Fig. 1d, Extended Data Fig. 4a and Methods), each dot in the 3D model represents the cell population at the specific positional address. The ‘3D corn plot’ model was utilized as basis for spatial mapping of single cells (see details in the next section).

**Figure 4.**
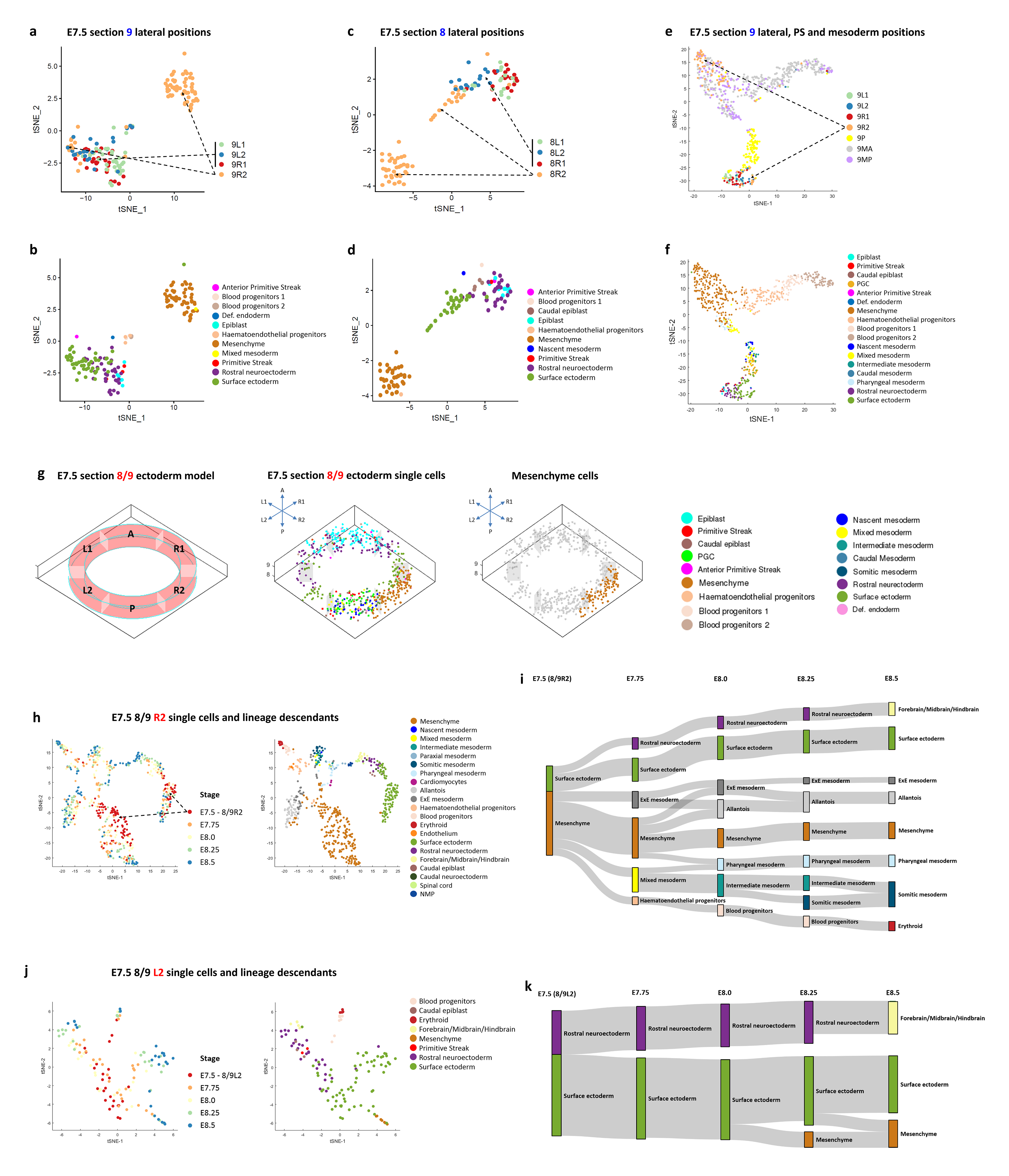
Heterogeneity of cell types and developmental trajectories of single cells in the proximal-lateral ectoderm of E7.5 embryo. **a-f.** *t*-SNE plots showing the heterogeneous cell clusters in position-9R2 and −8R2. Cells are annotated by the position (**a, c, e**) and cell-type (**b, d, f**). Dashed arrows (in **a, c, e**) denote the heterogeneous cell clusters in position-9R2 and −8R2, versus other lateral positions. Panel **a**-**d** showed the single cells of proximal-lateral ectoderm positions in section-9 (**a**, **b**) and −8 (**c**, **d**). Panel **e**, **f** showed the single cells of proximal-lateral ectoderm, primitive streak and mesoderm positions in section-9. **g**. The spatial distribution of single cells in the Annulus Model of ectoderm at Geo-seq section-8/9. **h-k.** *t*-SNE plots (**h, j**) and the molecular trajectories (**i, k**) of single cells (imputed using the Population Tracing algorithm) at position-8R2/9R2 (**h, i**) and position-8L2/9L2 (**j**, **k**) in E7.5-E8.5 embryos. Developmental timepoints (stage) and cell types (see legend) are indicated in the *t*-SNE plots. Cell types in E7.75-E8.5 embryos are annotated according to the ‘Gastrulation Atlas’.

## A single-cell resolution 3D molecular atlas

Single-cell RNA-sequencing (scRNA-seq) approaches have been used to profile the molecular feature of individual cells during early development^2, 8^. But most single-cell studies have profiled dissociated populations of cells, where spatial information of the single cells in the embryo is lost. Previous works of mapping the location of cells in biological structures on the basis of the concordance of the gene expression profile^10, 14^ have been confounded by mathematical uncertainties and false-positives^15^. A high-value attribute of the spatio-temporal transcriptome is the amenability of mining the dataset to identify population-specific signature transcripts as zipcodes (Fig. 1a) that could be applied for imputing the position of single cells in the germ layers of E6.5- E7.5 embryos^10, 16^. Leveraging our spatio-temporal molecular atlas and single-cell RNA-sequencing (scRNA-seq) dataset, we developed an analytics methodology that comprises tiered algorithms to infer the spatial distribution of single cells and to reconstruct a single-cell resolution 3D molecular atlas (Fig. 1e).

In our mapping study, the 3D corn plot model was used for mapping the spatial coordinates of single cells. By employing a Multi-Dimension Single-Cell Mapping (MDSC Mapping) algorithm (Fig. 2a and Methods), single cells were mapped by the zipcodes embedded in their transcriptome to the inferred position in the model. To evaluate the level of precision of positional mapping, we tested the mapping of single cells isolated from known positions at five developmental timepoints between E6.5 and E7.5. The results showed that the single cells could be mapped to their best-fit site of origin at significant confidence-level (PCC values at 0.74-0.97, Fig. 2b, Extended Data Fig. 4b and Methods). Other technologies, such as MERFISH and MERSCOPE, are amenable for capturing spatial transcriptome at single cell resolution, but have limited gene detection rate^15^ compared with Geo-seq, and therefore are less suitable for the spatial mapping of single cells.

Drawing from the single-cell transcriptome data of mouse embryo at gastrulation to early organogenesis (‘Gastrulation atlas’, Fig. 2c, d)^2^, we first applied MDSC Mapping algorithm and mapped the 25 embryonic cell types to the germ layers of the E6.5-E7.5 embryos (Fig. 2e and Extended Data Fig. 4c-l). To visualize the spatial distribution of the cells in the germ layers, an Annulus Model was applied to display “Geo-seq position” of single cells (Fig. 1e, Extended Data Fig. 5a-e and Methods) in a series of 3D-rendered spatio-temporal maps. After mapping single cells to the best-fit position in the Annulus Model (Extended Data Fig. 5d, e), we applied mathematical modelling based on Bubble Sort Algorithm to re-position single cells within each Geo-seq position by incorporating information of vectorially graded molecular activity (e.g., gradient of signaling activity and gene expression levels) (Fig. 1e, Extended Data Fig. 5f and Methods) that is known to associate with the regionalization of cell fate. This mathematical model enables an effective imputation of the coordinates of single cells in the Geo-seq zipcode position (Fig. 2f-h). By taking into consideration of the pattern of distribution across the developmental stage, the data could be rendered digitally into the Spatio-Temporal (4D) Atlas at single-cell resolution (Fig. 2f and Extended Data Fig. 6a). This 4D Atlas provides unique insights into the spatial distribution of, for example, specific group of single cells in the germ layers of E6.5-E7.5 embryos (Fig. 2g and Extended Data Fig. 6a) and, in specific cases, the *Pou3f1*-expressing cells in the epiblast/ectoderm (Fig. 2h), and *T*-expressing cells in the three germ layers (Extended Data Fig. 6b) during gastrulation.

**Figure 5.**
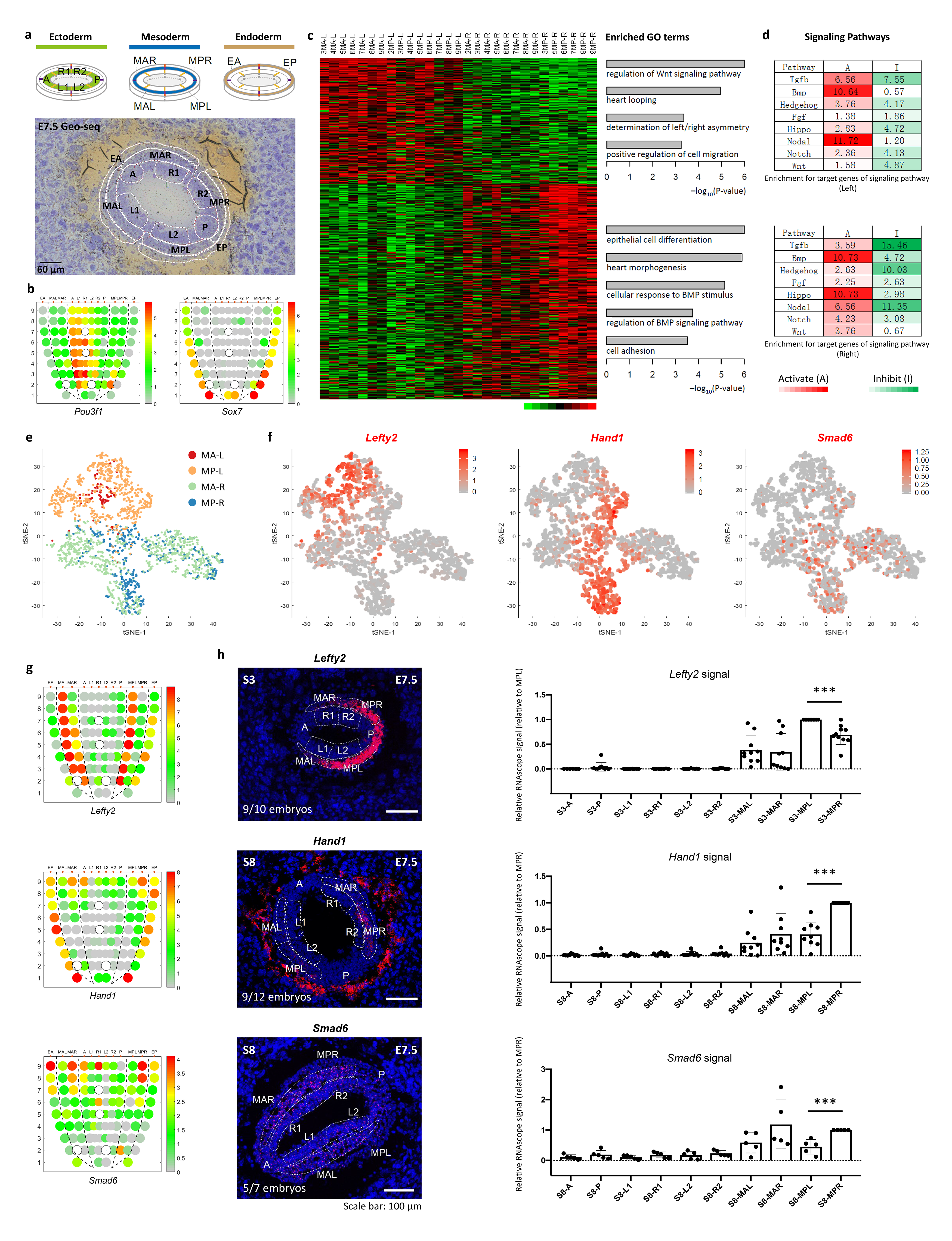
Left-right asymmetry at late-gastrulation stage. **a.** The strategy of laser capture microdissection of cell samples of E7.5 embryos. For the ectoderm and endoderm germ layers, the same Geo-seq strategy was applied as in Extended Data Fig. 1. The mesoderm germ layer was partitioned into MAL (anterior left mesoderm), MAR (anterior right mesoderm), MPL (posterior left mesoderm) and MPR (posterior right mesoderm) areas for sampling. Sampling areas are shown in histology images; scale bar, 60 μm. **b.** Corn plots showing the spatial pattern of expression of *Pou3f1* and *Sox7*. Hollow circles indicates missing samples. **c.** Heat map showing the differentially expressed genes (DEGs) of the left lateral mesoderm (n = 518) and right lateral mesoderm (n = 881) (p < 0.05, fold change > 1.5). The enriched gene ontology (GO) terms for each group were listed on the right (p < 0.01). **d.** The enrichment for target and response genes of development-related signaling pathways in the left and right mesoderm. Signaling activity: red, activating (A); green, inhibitory (I). The significance of –log_10_(FDR) value in each cell was calculated by one-sided Fisher’s exact test followed by Benjamini-Hochberg correction. **e.** Deconvolution analysis inferred the proportion of left and right lateral mesoderm cell populations, and visualized on t-SNE plot. Cells are colored by inferred positions: MA-L, MP-L, MA-R and MP-R. **f.** *t*-SNE plots showing the distribution of *Lefty2*-expressing cells in the left mesoderm, and *Hand1-* and *Smad6-*expressing cells in the right mesoderm. **g.** Corn plots showing the distribution of *Lefty2*-expressing cells in the left mesoderm, and *Hand1-* and *Smad6-*expressing cells in the right mesoderm. **h.** RNAscope analysis validated the bilaterally asymmetric expression of *Lefty2*, *Hand1* and *Smad6* in the selected transverse sections (S-numbered, reference: Fig 5g). Right panels summarize the quantified signal intensity and statistical results, n=9 for *Lefty2* and *Hand1,* n=5 for *Smad6*.

**Figure 6.**
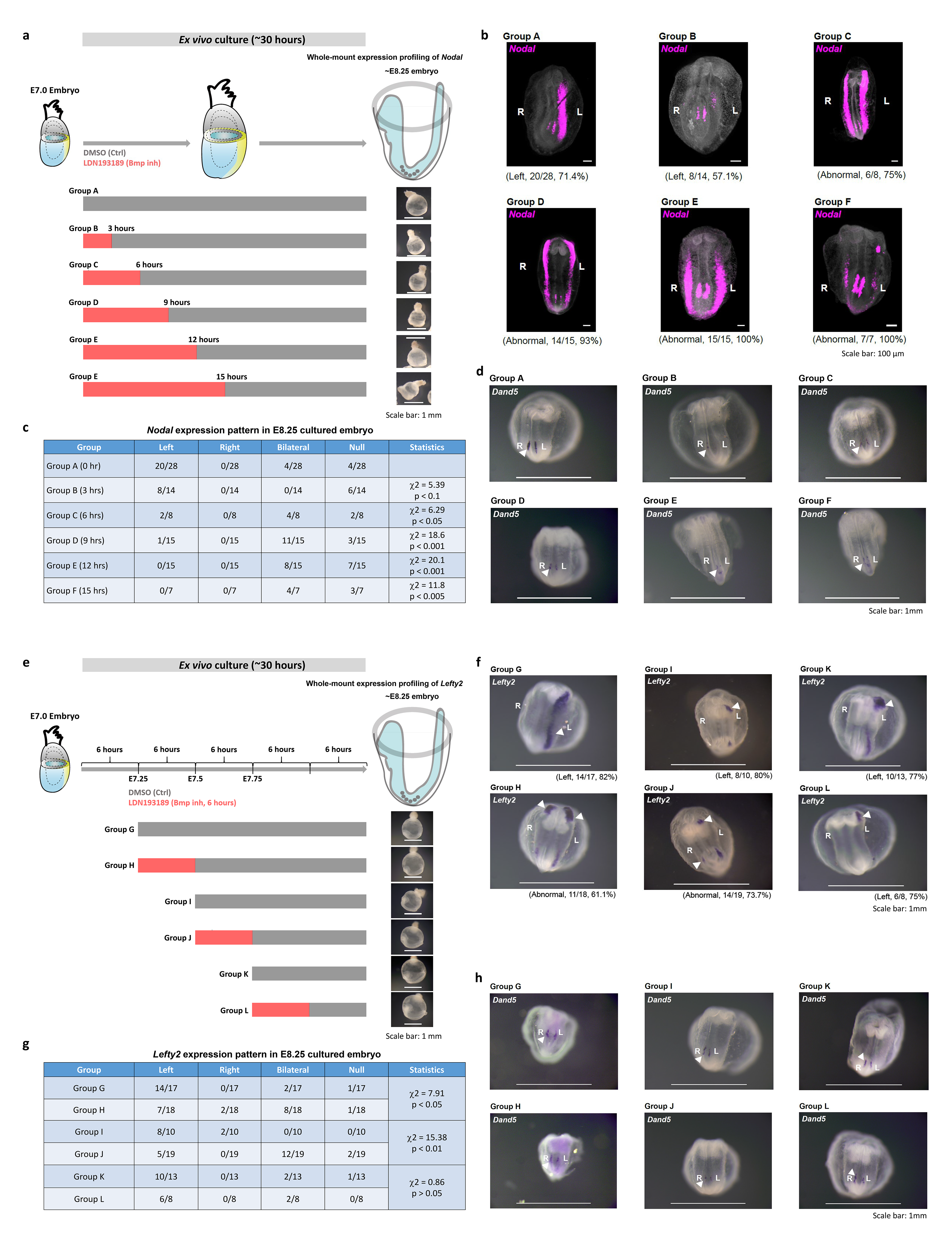
The temporal roles of gastrulation stage BMP signaling pathway in regulating left-right asymmetry. **a.** Experimental strategy of *ex vivo* culture and analysis of embryos following chemical inhibition of BMP activity for 3, 6, 9, 12 and 15 hours beginning on E7.0. **b.** Whole-mount RNAscope analysis of embryos after 30 hours of *ex vivo* culture, showing the expression of *Nodal* in the lateral mesoderm and the node (ventral view). The frequency of left-sided expression of *Nodal* is shown for Group A (untreated control) and Group B (3 hours treatment), and the frequency of abnormal (bilateral) expression of *Nodal* is shown for Groups C (6 hours treatment), D (9 hours treatment), E (12 hours treatment) and F (15 hours treatment). **c.** Table showing the pattern of *Nodal* expression in the cultured embryos collected at 30 hours in vitro (equivalent to E8.25). **d.** Whole-mount in situ hybridization of *Dand5*/*Cerl2* in cultured embryos from indicated experimental groups. **e.** Experimental strategy of *ex vivo* culture and analysis of embryos following chemical inhibition of BMP activity for 6 hours beginning at E7.25 (Group G and H), E7.5 (Group I and J), and E7.75 (Group K and L) stages. **f.** Whole-mount in situ hybridization of *Lefty2* after *ex vivo* culture (equivalent to E8.25), showing the expression of *Lefty2* in the lateral mesoderm (ventral view). The frequency of annotated pattern of *Lefty2* is also shown for each group. **g.** Table showing the pattern of *Lefty2* expression in the cultured embryos collected at equivalent E8.25 stages. **h.** Whole-mount in situ hybridization of *Dand5*/*Cerl2* in cultured embryos from indicated experimental groups.

The single-cell mapping results were generally consistent with prior knowledge of the identity of cell types annotated based on expression markers and lineage inference^1, 2^. For example, the anterior primitive streak cells were mapped to the distal posterior epiblast (positions 1P-4P) at E7.0 (Extended Data Fig. 4h), and rostral neuroectoderm cells were mapped to anterior regions of the ectoderm of E7.5 embryo (Extended Data Fig. 4l). However, some cell types were mapped to areas of the germ layer that are apparently inconsistent with their cell identity. For example, cells of the primitive streak were mapped widely in the epiblast outside the primitive streak (Extended Data Fig. 4d, f, h, j, l) and cells that are annotated as nascent mesoderm were mapped predominantly to the posterior epiblast (6P-11P), besides the mesoderm layer at E7.0 (Extended Data Fig. 4h). It might be that these cells in the epiblast represent a transitional cell state and not the fully specified cell type^3, 17^. This observation raised the possibility that the primitive streak cells and nascent mesoderm mapped to the epiblast may represent a population of progenitor cells at the transition of their allocation from epiblast to the emerging germ layers. Drawing on the knowledge of the prospective fate of cells in the epiblast of gastrulating embryos^1, 18^, we re-annotated the single cells mapped to the epiblast as progenitor/precursor cells (the intermediate cell types) of the germ layer derivatives based on the findings of heterogeneity of cell populations at a coordinate position in the germ layers (see next section).

## Heterogeneity of space-registered cell population

To visualize the composition of single-cell population at each Geo-seq position, an optimization algorithm based on Euclidean distance was applied (Fig. 3a and Methods). Using the gene-expression matrix of single cells mapped to a specific position as an input, computing a normalized Euclidean distance matrix followed by applying the Least Square Method, an optimized coordinate was assigned to each single cell. Applying this optimization algorithm, we simulated the spatial pattern of cell types within each Geo-seq position. For example, for position-6P at E6.75, this refined mapping revealed the presence of a heterogeneous cell population of the epiblast, primitive streak and nascent mesoderm cells (Fig. 3b, c). This finding prompted us to examine the heterogeneity of cell types in all the Geo-seq positions in the germ layers of the embryo at the five developmental timepoints.

To uncover the heterogeneity of single-cell population in the Geo-seq position, we applied t-SNE to re-cluster cells at each position in the germ layers of embryos at the five developmental stages (Extended Data Fig. 7). This analysis revealed that some cell types that were previously annotated differently were transcriptionally similar, illustrating the challenges of grouping of cells into discrete entities by transcriptome primarily. For these cells with different annotations that mapped to an unexpected germ-layer domain but displayed proximity of transcriptome, we re-annotated the cell identity based on both marker gene expression and prior knowledge of the prospective cell fate gleaned from fate-mapping and lineage tracing studies^5, 6^ (Extended Data Fig. 7 and 8). For example, analysis of the heterogeneity of cell types identified two clusters of single cells at Geo-seq position-6P in the posterior epiblast of E6.75 embryo (Extended Data Fig. 9a-c). One cluster displayed upregulation of mesoderm genes and enriched gene ontology (GO) terms of mesoderm development, and the other cluster (Cluster 2) displayed transcription signature of ectoderm. These two clusters were therefore annotated as ‘PS → Mes’ and ‘Epi→Ect’ respectively (Extended Data Fig. 9c, see methods for nomenclature). RNAscope analyses validated the expression of *Mesp1* (Extended Data Fig. 9d, e) and co-existence of *Pou3f1*-positive cells as well as *T*- positive cells in the proximal-posterior epiblast of E6.75 embryo (Extended Data Fig. 9d, f). On the basis of the fraction of re-annotated cell types of every Geo-seq position, a series of Heterogeneity Map was constructed for the gastrula embryo (Fig. 3d), and the presence of all cell types in the three germ layers during gastrulation was identified (Fig. 3d and Extended Data Fig. 8).

**Figure. 7.**
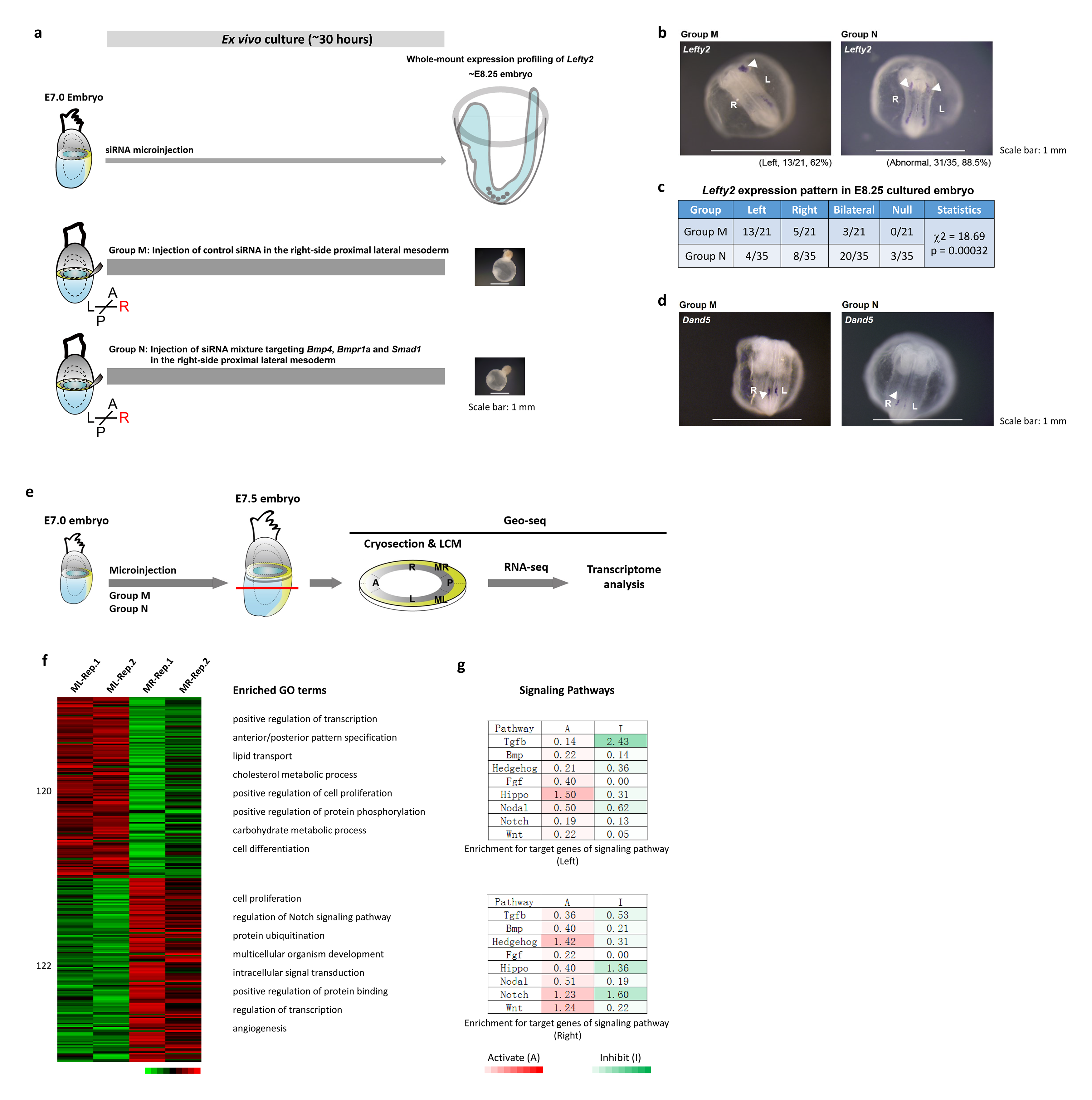
The spatial roles of gastrulation stage BMP signaling pathway in regulating left-right asymmetry. **a.** Experimental strategy of *ex vivo* culture and analysis of embryos following siRNA microinjection beginning at E7.0 stage. Group M: Injection of control siRNA; Group N: Injection of Bmp siRNA mixture. **b.** Whole-mount in situ hybridization of *Lefty2* after *ex vivo* culture (equivalent to E8.25), showing the expression of *Lefty2* in the lateral mesoderm (ventral view). The frequency of annotated pattern of *Lefty2* is also shown for each group. **c.** Table showing the pattern of *Lefty2* expression in the cultured embryos collected at equivalent E8.25 stages. **d.** Whole-mount in situ hybridization of *Dand5*/*Cerl2* in cultured embryos (equivalent to E8.25) from indicated experimental groups. **e.** Schematic diagram showing the workflow of GEO-seq for microinjected embryos. **f.** Heatmap showing the DEGs of the proximal-left mesoderm (n = 120) and proximal-right mesoderm (n = 122) (p < 0.01, fold change > 1.5) in the siRNA-KD embryos. Replicates: Rep-1, Rep-2. The enriched gene ontology (GO) terms for each group were listed on the right. **g.** The enrichment for target/response genes of development-related signaling pathways in the proximal-left and proximal-right mesoderm of the BMP siRNA microinjected embryos. Signaling activity: red, activating (A); green, inhibitory (I). The significance of –log_10_(FDR) value in each cell was calculated by one-sided Fisher’s exact test followed by Benjamini-Hochberg correction.

## Inferring the Developmental Trajectory

The information of cellular heterogeneity further revealed the progressive diversification of cell types in the germ layers (Fig. 3d), which might mirror the events of lineage development during gastrulation. The maps showed that a homogeneous cell population is found in epiblast and ectoderm domains initially and the heterogeneity in cell type composition emerges during gastrulation in cell populations in the posterior epiblast/ectoderm, the primitive streak, the mesoderm and the endoderm. To reveal the developmental connectivity of the cell populations with increasing heterogeneity, the Population Tracing algorithm (Extended Data Fig. 2f and Methods) was applied to infer the putative molecular trajectory of the primitive streak like cells (PSLCs) in the posterior epiblast of E6.5 embryo to cells in the germ layers at advancing stages of gastrulation (Fig. 3d). Embedded in the inferred trajectories is the information of the regulatory networks that drive lineage development, for example, the derivation of PS-Mes cells from E6.5 PSLCs, and subsequent allocation of PS-Mes cells to diverse mesodermal cell types at E7.5 (Fig. 3e). It is worth noting that the PSLCs diverge to progenitors of three germ layers at E6.75. A portion of cells in the proximal-posterior epiblast contributes to epiblast/ectoderm lineage, while the bulk of PSLCs is partitioned into proximal and distal cell groups that were allocated to mesoderm and mesendoderm lineages. The ‘PS → Mes’ cells from proximal primitive streak contribute different mesoderm derivatives at later stages, including the blood and cardiac mesoderm that emerge from proximal primitive streak-mesoderm populations at E7.0. In contrast, the PSLCs in the distal primitive streak are inferred to give rise to the mesendoderm progenitor by E7.25. These findings further showed that regionalization of cell types that underpin tissue patterning in the germ layers is evident at the molecular level as early as the mid-gastrula stage. In our study, we identified the putative mesendoderm progenitor (MEP) in the E7.25 embryo, which may contribute to both mesodermal and endodermal lineages (Fig. 3e and Extended Data Fig. 10a, b). The MEP was not identified in another single-cell study^19^, leading to the inference that the bifurcation of mesoderm and endoderm lineage occurs rapidly on exit from the pluripotent state, such that no self-renewing, albeit transiently, mesendodermal progenitor cells can be defined. Furthermore, cells that were annotated as ‘primitive streak’ were highly correlated with the PSLCs in our study. Overall, the inferred molecular trajectories are broadly in line with the prospective cell fate of the epiblast cells, which provide an entry point for imputing molecular control of the specification and differentiation of cell types derived from the primitive streak like cells at successive timepoints of germ layer development.

## Chirality of Cell Populations in Proximal Epiblast

An unexpected discovery of the cellular heterogeneity of the epiblast was the finding of different cellular composition of the proximal-lateral epiblast located on contralateral sides (positions 8R2 and 9R2 versus 8L2 and 9L2) of the E7.5 embryo. The populations on the right side are an admixture of cell types while those on the left side are homogeneous (Fig. 3d, e). To delineate the diversity of cell types in the population, we computationally re-clustered the single cells mapped to position-8R2 and −9R2 and the adjacent positions: 8R1, 9R1 and contralateral positions 8L1, 8L2, 9L1 and 9L2 in the proximal-lateral epiblast, as well as position-9P in the primitive streak and position-9MA and −9MP in the mesoderm. The clustering analysis affirmed the heterogeneity of cell types in position 8R2 and 9R2 (Fig. 4a-d and Extended Data Fig. 11a-d) – in particular, a large fraction of the cells at these two positions were allocated to a distinct cluster that was enriched for mesenchyme characteristics^2^, in contrast to the remaining cells, which grouped with cells from other positions and were enriched for ectoderm characteristics. The mesenchyme cell type is transcriptionally similar to cells in the mesoderm and is tightly clustered with cells in position-9MA and −9MP (Fig. 4e, f and Extended Data Fig. 11e, f). Applying the Euclidean distance-based Optimization Algorithm, we recapitulated the spatial distribution of single cells within ectoderm section-8/9 (Fig. 4g), where mesenchyme-like cells, distinct from other cell types, were enriched in position-8/9R2. To identify the descendants of these cell clusters in position-8R2 and −9R2, we applied the Population Tracing algorithm to infer the descendants of these cells in E7.75-E8.5 embryos (based on single-cell data in the ‘Gastrulation Atlas’)^2^ (Extended Data Fig. 11g). The cell clusters follow different trajectories, with the mesenchyme cluster giving rise to a multitude of mesodermal tissues, and the ectoderm cluster to the surface ectoderm and rostral neuroectoderm (Fig. 4h, i and Extended Data Fig. 11h). These two cell clusters at position 8/9R2 are likely to be the progenitors of the lateral mesoderm, and the surface ectoderm and neuroectoderm respectively. In contrast, cells that populate other six proximal-lateral positions (8/9L1, 8/9L2 and 8/9R1) contribute primarily to ectoderm derivatives and make a minor contribution to ectomesenchyme (presumptively the neural crest cells) (Fig. 4j, k and Extended Data Fig. 11i-l). Consistent with previous findings^1^, anterior lateral epiblast (L1, R1) contributes mainly to the neuroectoderm lineage, whereas posterior lateral epiblast (L2) preponderantly contributes descendant to the surface ectoderm. This preponderant contribution of the posterior proximal epiblast on right side of the embryo to the mesoderm derivatives may implicate an alignment of this developmental event to the specification of left-right (L-R) body asymmetry of the embryo.

## Initiation of Laterality in the Body Plan

To examine the L-R asymmetry of the germ layers in more depth, we performed Geo-seq analysis focusing on the domain of molecular activity in the mesoderm of E7.5 embryos (Fig. 5a, b and Extended Data Fig. 12a-f). In addition to the L-R difference in the proximal ectoderm (Extended Data Fig. 13a, c-e and h), we noticed that the proximal R2-related genes were more divergent in the right-side proximal lateral mesoderm (Extended Data Fig. 13b, f, g and i). Analyses of the spatial transcriptome revealed that the mesoderm cell population on contralateral sides displayed different profiles of gene expression and enrichment of functional gene ontology (Fig. 5c and Extended Data Fig. 14a). Analysis of whole-population transcriptome revealed left versus right differences in the enrichment of signaling activity of TGFβ, BMP, Hippo and Nodal pathway (Fig. 5d). By deconvolution analysis using the spatial transcriptome data, single mesoderm cells were allocated to the four domains (MA-L, MP-L, MA-R, MP-R) in the mesoderm layer (Fig. 5e, f and Extended Data Fig. 14b). Analysis of the transcriptome of the single cell populations revealed that the proximal mesoderm population (sections 7-9) on contralateral sides of the embryo displayed different gene expression profiles (Extended Data Fig. 14c) and the BMP and Hippo pathway activity was enriched in the mesoderm on the right side of the embryo (Extended Data Fig. 14d). Lateral mesoderm populations on contralateral sides were transcriptionally distinct. The side-specific enhanced pattern of marker genes (left: *Lefty2*, right: *Hand1* and *Smad6*) was validated by RNAscope and RT-qPCR analyses (Fig. 5g, h and Extended Data Fig. 14e, f).

Left-right (L-R) asymmetry of the body plan is manifested at the early-organogenesis stages (E8.25-E8.5), by the looping of heart and rotation of the epithelium of foregut portal^20, 21^, and the asymmetric Nodal signaling activity and left-sided *Lefty2* expression in the lateral mesoderm (Extended Data Fig. 14g)^22, 23^. In the mouse, the L-R tissue patterning is reputed to be initiated by asymmetric Nodal signaling activity in the node (the left-right organizer) at the post-gastrulation stage (∼E7.75), coupling with the propagation of signaling activity to the lateral plate mesoderm (LPM) to activate *Nodal* and Nodal target genes by the early-somite stage (∼E8.25, Extended Data Fig. 14g)^22, 24^. BMP signaling has been known to modulate the Nodal signaling activity and the downstream signal transduction activity. During the establishment of laterality at the early-somite stage, symmetric BMP activity in the lateral plate mesoderm sets a bilateral repressive threshold for Nodal/Smad4-dependent *Nodal* activation. Super-activating the BMP activity by overexpressing constitutive ALK6 receptor in the mesoderm on the left side of the post-gastrulation mouse embryo can counteract the Nodal/Smad4 activity and leads to disruption of the asymmetric Nodal downstream gene activity in the lateral plate mesoderm^23^. To test the impact of asymmetric BMP signaling activity in the mesoderm at gastrulation on the establishment of body laterality, mid-streak embryos (E7.0) were treated *ex vivo* with BMP inhibitor (LDN193189) for a duration of 3 to 15 hours followed by Geo-seq analysis (at 12 hr *ex vivo* = E7.5 late-streak stage) or RNA in situ hybridization (at 30 hr *ex vivo* = E8.25 early-somite stage) (Fig. 6a and Extended Data Fig. 15a). Early-somite stage embryos that were treated with inhibitor for more than 3 hours from E7.0 showed abnormal pattern of *Nodal* expression in the lateral mesoderm (Fig. 6a-c), with a reduced frequency in the left-sided expression, and an increased incidence of bilateral and absent expression (6 hours: 75%, 9 hours: 93%, 12 hours: 100%, compared to control: 28%) (Fig. 6b and c). In the treated embryos (Group D: 9 hours treatment) at the stage equivalent to E7.5, Geo-seq analysis of the transcriptome revealed that BMP pathway activity was down-regulated in the right-side proximal lateral mesoderm while remained unchanged on the left-ride (Extended Data Fig. 15b) and the asymmetric pattern of gene expression in the lateral mesoderm has diminished (Extended Data Fig. 15c, d). The disruption of L-R asymmetry of Nodal activity in BMP-inhibited embryo raises the possibility that specification of the laterality of the body plan is initiated at late-gastrulation stage, ahead of the formation and functionalization of the L-R organizer. It was noted that in the BMP-inhibited embryos, *Nodal* and *Dand5* remained expressed in the peri-node tissue (Fig. 6b and d), suggesting that the action of BMP signaling in L-R patterning may be independent of the function of the L-R organizer. To delineate the stage-specific impact of BMP signaling on L-R patterning, the embryos were treated, beginning at different stages (E7.25, E7.5 and E7.75), for 6 hours by LDN193189 inhibition (Fig. 6e). Blocking BMP activity at the mid-late streak stage (E7.25) and the late-streak stage (E7.5), but not at the early-head-plate stage (E7.75), led to disruption of the asymmetric *Lefty2* expression in the lateral plate mesoderm of the early-somite stage embryo (Fig. 6f and g), while *Dand5* expression at the node region was not consistently changed (Fig. 6h). These findings point to a stage-specific requirement of asymmetric BMP activity during late gastrulation for establishing L-R molecular asymmetry.

To elucidate whether the enhanced BMP activity on the right side of the gastrulating embryo is critical for the acquisition of L-R molecular asymmetry, E7.0 embryos were treated by combined siRNA knockdown (KD) of *Bmp4*, *Bmpr1a*, and *Smad1* in the right lateral mesoderm (Fig. 7a and Extended Data Fig. 15e). The siRNA-KD embryos cultured to the early-somite stage displayed increased incidence of bilateral, ectopic and no expression of *Lefty2* in the lateral plate mesoderm (Fig. 7b-c), but proper node formation (Fig. 7d). Geo-seq analysis revealed that the early asymmetry, which maintained in control embryos, were abolished after siRNA KD (Fig. 7e and Extended Data Fig. 15f, g). Moreover, differentially expressed genes (DEGs) analysis of left versus right mesoderm of the siRNA-KD embryos identified less DEGs than in control embryos and revealed no significant difference of BMP signaling enrichment (Fig. 7f and g). In comparison, embryos that were subject to siRNA KD in the left lateral mesoderm maintained asymmetric *Lefty2* expression (Extended Data Fig. 15h-j). These results point to a critical requirement of the enhanced BMP activity on the right side of the embryo at late gastrulation stage for the establishment of laterality of the body plan.

To further elucidate whether the BMP signaling in the gastrulating embryo has mechanistic connectivity to the function of L-R organizer^25^ in mediating ciliary nodal flow and signal transduction, we assessed the impact of loss of *Pkd1l1*, *Cerl2* and *Dnah11* function^26–29^ on the asymmetric BMP signaling activity in the embryo. At the early-somite-stage (E8.25-E8.5), all three loss-of-function mutants displayed disruption of the asymmetric pattern of *Lefty2* expression (Extended Data Fig. 16a, b, d, e, g, h, i). However, the asymmetric expression pattern of *Lefty2* in left mesoderm and *Hand1* in right mesoderm at late gastrulation stage (E7.5) remained unchanged (Extended Data Fig. 16c, f, j and Extended Data Fig. 17a-c). Modulation of BMP signaling activity is known to counteract Nodal signaling activity in eliciting left-right asymmetry in the lateral mesoderm^23^ at more advanced stage of development. Thus, enhancement of BMP signaling activity in the mesoderm on the right side of the body at late gastrulation may represent an early symmetry-breaking event which is independent of the formation and function of the L-R organizer.

## Discussion

This time and space-resolved single-cell molecular atlas of mouse gastrulation adds to the knowledge base of the developmental trajectory of cell populations in the germ layers and molecular networks of cell-fate decisions during this critical period of early embryo development. This transcriptome resource has enable the charting of two developmental events in germ layer development: changes in the heterogeneity of cell types in position-registered cell populations, and the molecular trajectories of PS/PSLCs to germ layer derivatives. Combining Geo-seq, scRNA-seq and computational modeling, we uncovered that asymmetric BMP signaling activity in the mesoderm of embryo at late gastrulation may play a role in initiating L-R body patterning. BMP signaling activity is known to modulate Nodal signaling activity in the lateral mesoderm during the specification of left-right asymmetry^23^. The enhanced BMP activity at late gastrulation may constrain Nodal signaling activity on the right side of the embryo and thereby predisposing the contralateral side for perceiving and responding to Nodal activity emanating from the L-R organizer. Whether the BMP activity in the mesoderm may influence the mesoderm-lineage propensity of the proximal lateral epiblast on the right side of the gastrulating embryo, and if the enhanced contribution of the epiblast to the lateral mesoderm has a role in L-R patterning are not known at this juncture. Nevertheless, our findings have pointed to that specification of L-R asymmetry may be initiated at late gastrulation, ahead of the acquisition of functionality of the L-R organizer. This study of L-R patterning highlights the attribute of the spatio-temporal molecular atlas to advance our understanding of early mammalian development, which is an empowering resource for guiding future research on the molecular and cellular mechanism of lineage differentiation and tissue patterning in the post-implantation mouse embryos.

## Materials and Methods

### Sampling of mouse embryos for Geo-seq analysis

All animal experiments were performed in compliance with the guidelines of the Animal Ethical Committee of the CAS Center for Excellence in Molecular Cell Science, Chinese Academy of Sciences. C57BL/6J embryos were harvested from pregnant mice at Day 6.5, 6.75, 7.0, 7.25 and 7.5 of gestation (day of vaginal plug detection = Day E0.5). The embryos progression of gastrulation at the five developmental timepoints were staged by the proximal-distal span of the primitive streak and anterior-posterior span of mesoderm layer^30^. The staging was further confirmed by the spatial domain of *T-* and *Mixl*-expressing cell population in the posterior (P) samples of the embryo. Embryos were collected immediately after harvesting in OCT medium (Leica Microsystems, catalogue no. 020108926). Cryosections of the embryo were processed for laser capture dissection to collect cell samples from the germ layers as previously described^1^. Cell samples were processed for Geo-seq analysis^11^. The acquired cDNA was allocated for RT-qPCR analyses of gene expression, and library preparation for transcriptome profiling (Novozyme, TruePrep DNA Library Prep Kit V2 for Illumina, TD-503). Next generation sequencing was performed on the Illumina Hiseq 2500 or Novaseq platform (Berry Genomics).

### Whole-mount in situ hybridization

RNA whole-mount in situ hybridization of the gastrula were performed as previously reported^31^. Briefly, embryos were fixed in 4% paraformaldehyde (PFA; sigma; #P6148) overnight, rehydrated through the series of 75%, 50% and 25% methanol at room temperature, washed by 3 times in DPBS, treated with 10 mg/mL proteinase K (Life Technologies, # AM2548) in PBS for 8 min and post-fixed for 20 min in 4% paraformaldehyde. Riboprobes were synthesized by DIG RNA labeling kit and in vitro transcription kit (Roche Applied science, 11277073910; mMESSAGE mMACHINE T7 ULTRA KIT, AM1345; MEGAclear™ Transcription Clean-Up Kit, AM1902). Post-fixed embryos were incubated with approximately 1 μg/mL of digoxigenin-labeled RNA probe at 68°C overnight, washed and stained with anti-digoxigenin antibody to visualize the expression pattern of the gene transcripts hybridized with the riboprobes.

### RNAscope

OCT embedded E7.5 mouse embryos were cryo-sectioned at 20 μm thickness serially from the distal to proximal region and mounted on electrostatic glass slides. Analysis of gene expression was performed using RNAscope® Multiplex Fluorescent Reagent Kit v2 (Advanced Cell Diagnostics, 323100) following manufacturer’s instructions with minor modifications (proteinase plus treatment for 15 min instead of proteinase IV for 30 min) using probes supplied by Advanced Cell Diagnostics: mm-Pou3f1 (436421), mm-Mesp1-C3 (436281-C3), mm-T-C3 (423511-C3), mm-Lefty2-C2 (436291-C2), mm-Nodal-C3 (436321-C3), mm-Smad6-C4 (528041-C4), mm-Hand1 (429651). Images were acquired using Leica TCS SP8 STED system. The fluorescence intensity of each region was calculated using Image J software following standard steps (8-bit format transformation > Setting threshold for background removal > Signal measurement). In order to minimize signal variance between embryos and experimental batches, the integrated fluorescent intensity of each region was normalized by calculating the relative ratio to the corresponding region (e.g., relative ratio to MPL region for *Lefty2*, relative ratio to MPR for *Hand1* and *Smad6*). For each RNAscope probe, at least three biological replicates were examined and statistical significance was assessed by student’s t-test.

### Small interfering RNA (siRNA) perturbation of BMP signaling

siRNA oligos for *Bmp4*, *Bmpr1a* and *Smad1* tagged with Cy3 orange-fluorescent dye and fluorescein amidite (FAM)-labelled negative control siRNA were synthesized by GenePharm. The siRNAs were firstly dissolved using RNase free water to make 50 μM stock solutions following the manufacturer’s instructions. The efficiency of each oligo for knocking-down the transcripts were pre-determined in mouse embryonic stem cells. The siRNA with the highest knockdown efficiency were selected for the experiments. Equal volume of the oligos for *Bmp4*, *Bmpr1a* and *Smad1* were mixed into one siRNA preparation. For lipofectamine mediated transfection, 4 μL siRNA mixture, 1.5 μL LipoFectamine 2000 (Invitrogen, 11668-027), 4.5 μL OptiMEM were pre-mixed and incubated at room temperature for 5 min. E7.0 mid-streak stage mouse embryos were collected from the pregnant mice and transferred to drops of PB1 medium placed on the 37°C warm plate under the microscope. Approximate 0.05 μL siRNA-lipid mix were micro-injected into the extracellular space between the ectoderm and endoderm of the right-side of the recipient embryo using a flat-tip microinjection pipette^32^. Into control embryos, the same volume of negative control siRNA was injected. The injected embryos were cultured *ex vivo* as described below.

### *Ex vivo* culture of mouse embryo

Mid-primitive-streak (E7.0) embryos were cultured in medium of 50% CMRL (Gibco, 11530037) and 50% rat serum, supplemented with 1xGlutamax (Gibco, 35050061), 1xNEAA (Hyclone, SH30238.01) and Glucose (4 mg/mL) under 5% CO_2_ incubator at 37°C. 1 μM chemical inhibitor of BMP signaling (LDN193189, Selleck, S2618-2mg) was added to the culture medium in the respective experimental groups. The cultured embryos were collected at various timepoints (Figure 6, 7 and Extended Data Fig. 15) for whole-mount RNA in situ hybridization or GEO-seq. The whole-mount RNAscope experiments were performed following the published protocol^33^. Images were captured using LiTone XL system (Light Innovation Technology Limited). Embryos subjected to GEO-seq were fixed in paraformaldehyde for 30 min at 4°C, dehydrated in an alcohol series, embedded in OCT compound and cryo-sectioned serially. Cells were sampled by laser capture microdissection following the published protocol^11^. The acquired samples were treated with protease K (Invitrogen, AM2546) at 56°C for 30 min, and the lysate was precipitated in ethanol solution and prepared for RNA-sequencing as previous described^11^.

### Pre-processing of RNA-seq data

Sequencing quality of raw sequencing data was evaluated by FASTQC. Tophat2 v2.0.4 program^34^ was used to map raw reads to mm10 version of mouse genome with the following parameters: -g 1 -N 4 --read-edit-dist 4 --microexon-search -G annotation.GTF. Mapping ratio was calculated based on the number of mapped reads and total reads for each sample. We calculated fragment per kilobase per million (FPKM) as expression level using Cufflinks v2.0.2 with default parameters^35^. For each embryo, genes with FPKM > 1.0 in at least two samples across all samples were retained for further analysis. Finally, the expression levels were transformed to logarithmic space by using the log2 (FPKM+1).

### Identification of spatial domain and zipcodes

For E6.5 and E6.75 embryos, zipcodes were identified as follows: 1) applied an adaptive clustering algorithm, Bayesian Information Criterion-Super K means (BIC-SKmeans) algorithm^36^, to identify an optimum number of clusters that can best capture the variance in the data, these optimum clusters were defined as spatial domains, and 2) Identify inter-domain differentially expressed genes (DEGs) using RankProd^37^ with P value < 0.05 and fold change > 1.5. Top 50 DEGs of each spatial domain were denoted as zipcodes for E6.5 and E6.75 embryos.

For E7.0, E7.25 and E7.5 embryos, zipcodes were identified as follows: 1) Use ComBat^38^ to remove potential batch effects based on expression of all genes (log_2_- transformed), due to variation between samples of different sequencing conditions. 2) Use Python program v2.7.13 to calculate the variance of each expressed gene across all samples and select top 6,000 genes as highly variable genes, then perform principle component analysis (PCA) using FactoMineR^39^ package in R. The top 50 highest and 50 lowest PC loading genes from the most significant PCs (E7.0, PC1-4; E7.25, PC1- 5; E7.5, PC1-5), which showed inter-domain specific expression patterns, were denoted as zipcodes for E7.0, E7.25 and E7.5 embryos respectively. Heatmaps were generated using Cluster 3.0 and JavaTreeView^40^.

### Population Tracing algorithm

To trace the developmental trajectory of cell populations in different spatial domains, we developed a digital tracing algorithm with the following computational procedures:

1. Use the union of genes in zipcodes of pair of stages of interest as the input gene set,
2. calculate the Euclidean distance of any two domains from two embryos of adjacent stages (for example, the distance between *D_A1* and *D_B2*, *D-A1* denotes domain 1 at time point A, *D-B2* denotes domain 2 at time point B) (i),

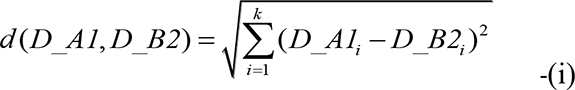
3. Calculate the mutual nearest neighbours (minimum matrix distance and variation less than 10%, the formula-(ii) below) for each domain from one developmental timepoint against the domains from the next developmental timepoint and connect the spatial domains. This procedure was repeated across the timepoints from the beginning to the end of the developmental series.

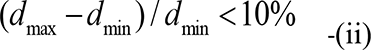

and 4) Convert the distance of any two connected spatial domains to logarithmic space by using log_2_ transformation and visualize the genealogy of cell populations by Sankey plot using Google Charts^1^.

### 3D Modeling for embryo structure and visualization of Geo-seq data

To mimic the spatial structure and visualize the spatial pattern of gene expression in the embryo, a geometric concentric-oval model that mirrored the architecture of the cup-shaped gastrula-stage mouse embryo was developed. The RNA-seq data of cell samples were assigned to the positions defined by spatial coordinates on the ‘3D corn plot’ model to depicted the spatial pattern of expression of the gene or gene group of interest, with the expression levels indicated by a color scale computed from the transcript counts in the RNA-seq dataset.

### Multi-Dimension Single-Cell Mapping (MDSC Mapping)

Based on the 3D model and the zipcode signature, a mathematical algorithm was developed for imputing the location of single cells in the germ layers of the mouse embryo. The mathematical operation included: 1) Calculate Spearman’s rank correlation coefficient (SRCC): The SRCCs between the expression values of the zipcodes of each single cell and all samples of the reference embryo were computed to generate, for example, 74 SRCC values for each single cell against 74 Geo-seq samples of E7.0 embryo, 2) Apply a spatial smoothing algorithm to determine the high-confidence location of each cell. This mapping method contrasts with the previous imputation method^10^ that single cells are mapped to the position of the maximum SRCC value. While the higher SRCC may indicate a strong probability of matching to a position, there were cases where a single cell could match to several adjoining positions. Therefore, for mapping the single cells, the top 3 matching positions with maximum SRCCs were extracted, then applied the formula (iii) below to calculate a 3D coordinate:

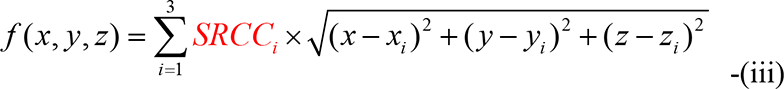

The distance between this location and every Geo-seq sample position was then calculated, the sample with minimum distance was determined as the best mapped position of the cell.

### Verification of the mapping efficiency

The efficiency of MDSC Mapping pipeline was evaluated by mapping single cells manually isolated from known positions in E6.5, E6.75, E7.0, E7.25 and E7.5 C57BL/6J embryos. First, embryos were dissected from the decidua in 10% FBS-DMEM medium, and cells were isolated from a specific position of the embryo by mouth pipetting guided by microscopy. Altogether, 19 single cells from E6.5, 32 single cells from E6.75, 37 single cells from E7.0, 24 single cells from E7.25 and 6 single cells from E7.5 were subjected to automatic Smart-seq2 amplification and library construction with the Agilent Bravo automatic liquid-handling platform^41^. Data preprocessing encompassed mapping, quality control and normalization using the same criteria previously established^1^. Based on the transcriptome data of these single cells, their position was mapped to the reference embryo by MDSC Mapping.

To access the accuracy of the mapping methodology, we performed the following analysis: 1) Apply image processing to identify the spatial positions of single cells isolated by pipetting, 2) Apply Gaussian Distribution to mathematically simulate the Confidence Intervals. For E6.5 - E7.25 embryos, we employ the standard normal distribution to simulate the confidence intervals:

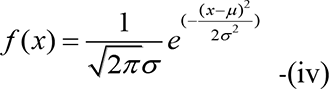

At E7.5, since the cell number is fewer, the standard normal distribution is adjusted to a more concentrated distribution: *f(x) ≈* 1. 3) Calculate the Pearson Correlation Coefficients (PCC) between the Confidence Intervals and MDSC Mapping results. Through this modeling protocol, we showed that the single cells could be mapped at significant fidelity to their site of origin, the PCC of MDSC mapping per stage is 0.7399 (E6.5), 0.8893 (E6.75), 0.7943 (E7.0), 0.7602 (E7.25) and 0.9738 (E7.5).

### Single cell datasets for positional mapping

The 10X Genomics single cell data were downloaded as raw files from the Gastrulation Atlas: https://github.com/MarioniLab/EmbryoTimecourse2018. Steps of quality control, normalization, batch correction and clustering were performed using the same criteria as previously described^2^.

### 3D modeling for single-cell resolution map

To visualize the spatial distribution pattern of single cells, an Annulus Model was developed to reconstruct the embryo spatial structure in single-cell resolution. The model comprises 1) Concentric annuli in the anatomical section of the embryo, from the inside outward, representing epiblast/ectoderm, mesoderm and endoderm germ layer respectively. 2) Division of the annulus into interior spaces matching the Geo-seq defined position. Single cells that mapped to a Geo-seq position, were distributed uniformly across the interior space of the position in each annulus section. 3) Within each interior space, apply Bubble Sort Algorithm to align the cells along the known gradient of gene expression level or signaling intensity. For example, on the basis that *Bmp4* expression in the posterior epiblast streak decreases along the proximal-distal axis of the embryo, the algorithm code (see below) was applied to rearrange cells in domain 8P in the E6.5 epiblast (x-cd, y-cd and z-cd are abbreviations of 3D spatial coordinate (x, y, z)), based on the gradient of *Bmp4* expression:

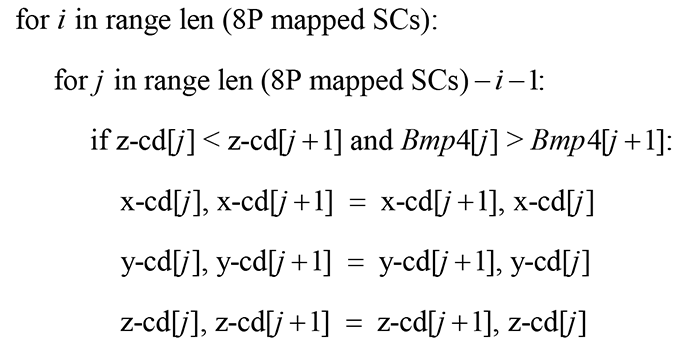

This imputation enabled the assignment of spatial coordinate specific for each single cell, and 4) Construct the 3D positional map for all single cells for the visualization of the spatial pattern of the single cells of different cell/tissue lineages that are individually defined by transcriptome features. Further information can be drawn from the 3D embryo map for the position of single cells displaying different level of expression (indicated by the color scale computed from the transcript counts in the ‘Gastrulation Atlas’) of gene/gene group of interest.

### The migrating mesoderm model

To model the mesoderm migration during gastrulation, we refined the Annulus Model by devising a Gradually Extended Annulus Model for the mesoderm of E7.0 and E7.25 embryos. From distal to proximal, the incomplete annuli that represent the anatomical section of the mesoderm progressively enveloping the epiblast from posterior to anterior. At E7.25, the annuli of proximal mesoderm (7-12M) completely encircles the epiblast. Bubble Sort Algorithm was then applied to assign coordinates to each single cell within interior space of each annulus of the mesoderm position.

### Euclidean-distance derived optimal coordinates

To refine the spatial distribution pattern of single cells within each Geo-seq position, an optimization algorithm based on Euclidean distance was developed. The mathematical operations include: 1) Compute the gene-expression matrix (GEM) of single cells mapped to a specific position, and transform the matrix to logarithmic space using log_2_((normalized count)+1). 2) Based on the logarithmic transformed matrix, calculate the Euclidean distance between every two cells to generate the Euclidean distance matrix (EDM). In order to adapt EDM to the interior space of Annulus Model, the EDM was normalized through the formula (iv) below to make the maximum matrix distance (*d_max_*) equivalent to the maximum length (*length_max_*) within the interior space.

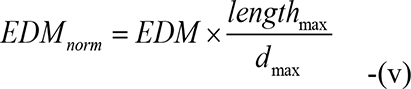

4) Apply the Least Square Method (v) under the constraint conditions of the spatial mathematical model of the interior space to derive the optimized coordinate for each single cell:

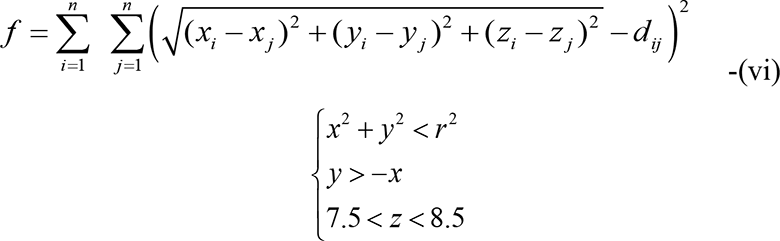

(This example shows the constraint conditions of the mathematical model for position 8P at E6.5 stage) and 5) Visualize the spatial pattern of single cells of different types in each position. All the spatial algorithms and 3D modelling were operated using MATLAB.

### Heterogeneity analysis

To reveal the composition of single-cell population mapped to a Geo-seq position, clustering analysis by *t*-SNE in Seurat^14^ was performed on the single cells for every position in the E6.5-E7.5 embryos. The cell clusters were annotated on the basis of marker gene expression and the knowledge of the prospective fate of cells in specific regions of the embryo, gleaned from lineage tracing and fate mapping studies^5, 6^. Based on the composition of annotated cell types in the single-cell population, a Heterogeneity Map was constructed for all the Geo-seq positions in the gastrula-stage embryos, with the cell-type composition displayed as pie charts in the corn plots. To construct the molecular trajectory of specific cell types across the developmental stages, single cells of same type or aligned with a specific lineage were grouped. The Population Tracing algorithm was applied using the averaged gene expression level data to infer the trajectories, which was visualized by Sankey plot using Google Charts^1^.

### Nomenclature for cell type annotation

The nomenclature ‘X→Y’ and ‘X(Y)’ represent different cell states. X represents the germ layer information of cell population. ‘X→Y’ indicates these cells are representing a transitional cell state from X to Y. And ‘X(Y)’ represents the precursor of a specified cell type (Y) in the germ layer X.

### Clustering and deconvolution of E7.5 mesoderm single cells

Transcriptome data of E7.5 mesoderm single cells were extracted from the ‘Gastrulation Atlas’. The Seurat package^14^ was used to perform single-cell clustering analysis and *t*-SNE was applied to visualize the results. To evaluate the representation of identified cell clusters in Geo-seq samples, we prepared cell-cluster labels for each single cell, the single-cell gene-expression matrix and all Geo-seq samples of mesoderm from the left and right side of the E7.5 embryo. CIBERSORT^42^ was applied to perform cell-cluster deconvolution analysis with default parameters (without quartile normalization). The proportion of each cell cluster in the population were visualized on *t*-SNE plot.

### Functional enrichment analysis

Functional enrichment of gene sets with different expression patterns was performed using the Database for Annotation, Visualization, and Integrated Discovery (DAVID)^43^ version 6.8.

### Signaling pathway enrichment analysis

In addition to BMP, FGF, Nodal, WNT, Notch and Hippo-Yap signaling pathways^1^, Hedgehog and TGFβ pathway were included in the analysis^44^. Potential signaling-target genes of each pathway were identified by comparing control samples with treatment samples using RankProd (P < 0.01 for Hippo-Yap and P < 0.001 for others) from published perturbation data (Gene Expression Omnibus accession numbers GSE48092, GSE41260, GSE17879, GSE69669, GSE15268, GSE31544, GSE58664 and GSE90567). Fisher’s exact test followed by Benjamin-Hochberg correction was applied to determine the significance of overlap of the target genes of signaling pathways in different DEGs groups.

### Generation of *Pkd1l1*, *Dand5* (*Cerl2*) and *Dnah11* (*iv*) mutant embryos

For the *Pkd1l1* rks mutant, substitution mutation in *Pkd1l1* gene^26^ was generated by adenosine base editing. *Cerl2* mutation was generated by introducing a stop codon in the first exon which phenocopied the Exon 2 deletion mutations^27^. For the *iv* mutant, the first P domain in the *Dnah11* gene was deleted^29^.

Small guided RNAs targeted to the respective genomic regions were designed by using the online tool Chop-chop (http://chopchop.cbu.uib.no/). The DNA fragments containing T7 promoter and sgRNA sequence and scaffold were transcribed *in vitro* using the MEGAshortscript Kit (Invitrogen, AM1354) and the products were purified using MEGAclear kit (Invitrogen, AM1908). DNA fragments for T7-CRISPR-Cas9^45^, T7-ABE 7.10^46^, and T7-YE1-BE4max^47^ were transcribed *in vitro* using MMESSAGE MMACHINE T7 Ultra Kit (Invitrogen, AM1345) and MEGAclear kit (Invitrogen, AM1908).

C57BL/6 female mice (4 weeks old) were superovulated and mated with the male C57BL/6 mice. 24 hours later, fertilized embryos were collected from oviducts. T7- CRISPR-Cas9 (for *Dnah11* mutant), T7-ABE 7.10 (for *Pkd1l1* rks mutant), T7-YE1- BE4max (for *Cerl2* mutant) mRNAs (100 ng/µl), and corresponding sgRNA (100 ng/µl) were mixed in HEPES-CZB medium containing 5 μg/ml cytochalasin B (CB) and injected into the cytoplasm of fertilized eggs using a FemtoJet microinjector (Eppendorf) with constant flow settings. The injected embryos were cultured in KSOM with amino acids at 37°C under 5% CO_2_ in air to reach the 2-cell stage after 24 hours *in vitro*. Two-cell embryos were transferred into pseudo-pregnant ICR female mice, and embryos were collected at E7.5 and E8.5 for analyses.

## Acknowledgements

The authors would like to thank F. Tang for critical discussions. This work was supported in part by the National Key Basic Research and Development Program of China (2019YFA0801402, 2018YFA0107200, 2018YFA0801402, 2018YFA0800100, 2018YFA0108000, 2017YFA0102700), the Strategic Priority Research Program of the Chinese Academy of Sciences (XDA16020501, XDA16020404), National Natural Science Foundation of China (31630043, 31900573, 31900454, 31871456), and China Postdoctoral Science Foundation Grant (2018M642106). P.P.L.T. was supported by the National Health and Medical Research Council of Australia (Research Fellowship grant 1110751).

## Author contribution

N.J., J.C.M. and R.W. conceived the study. N.J., J.C.M. and P.P.L.T. supervised the project. N.J., P.P.L.T. and X.Y. designed the experiments. X.Y., G.C., G.P. and Y.C. performed Geo-seq of embryos. Y.Q. performed animal husbandry. R.W., J.A.G. and J.C. analyzed the sequencing data. R.W., L.W. and J.C.M. generated the mathematical models. G.C. and X.Y. collected the region-specific single cells. X.Y. and Y.C. conducted the RNAscope and *ex vivo* culture experiments of mouse embryo. X.Y., L.Z., Y.C. and J.L. performed the siRNA perturbation experiments and established *Cerl2*, *Pkd1l1* and *Dnah11* mutant mice. R.W., X.Y., P.P.L.T., J.A.G. and N.J. wrote the paper with the help of all other authors.

## Competing interests

The authors declare no competing interests.

---

**Extended Data Figure 1.**
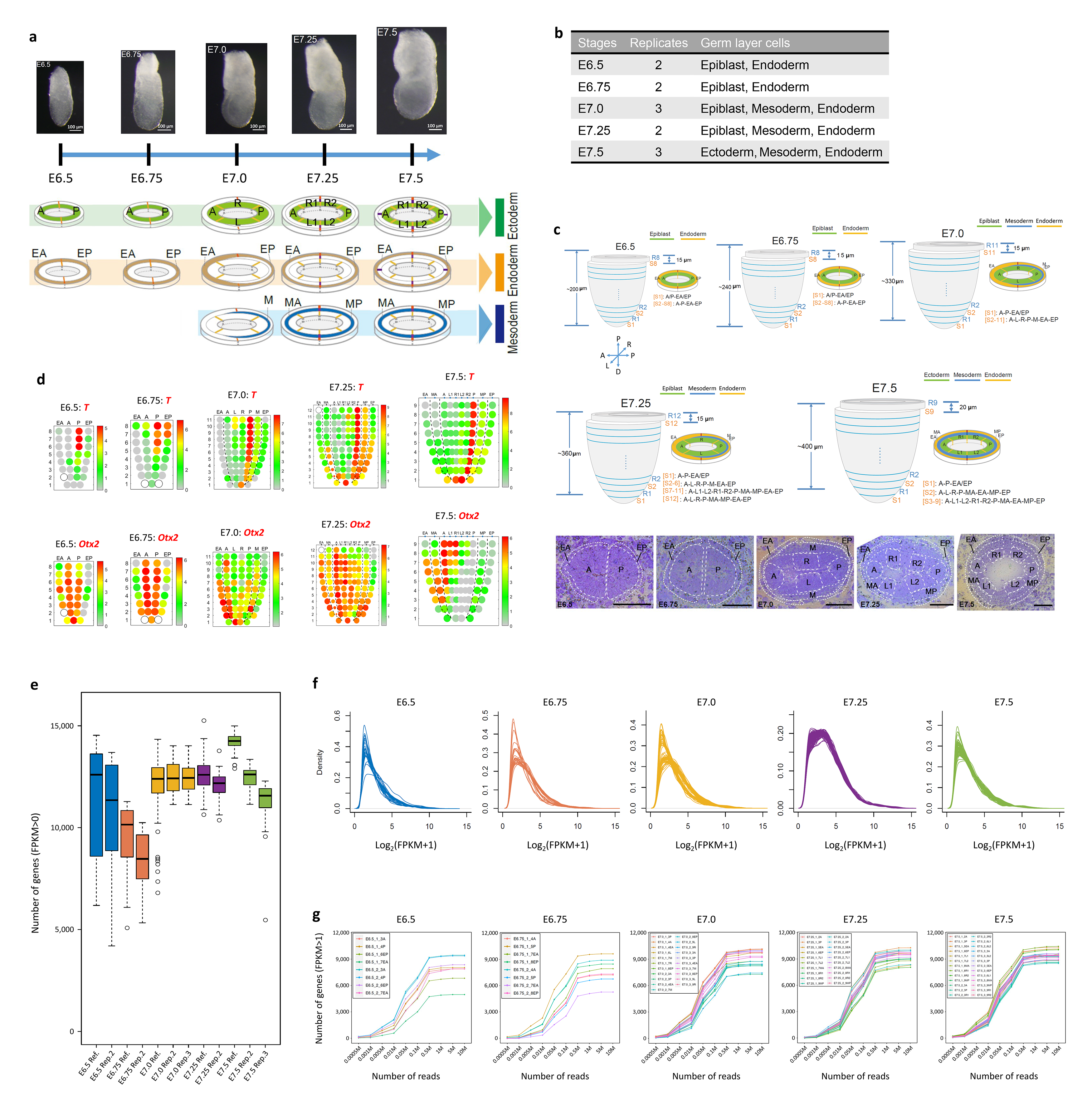
Geo-seq analysis. **a.** Schematics of laser capture microdissection of cell samples in E6.5-E7.5 embryos. A, anterior; P, posterior; L, left lateral; R, right lateral; L1, anterior left lateral; R1, anterior right lateral; L2, posterior left lateral; R2, posterior right lateral; M, mesoderm; MA, anterior mesoderm; MP, posterior mesoderm; EA, anterior endoderm; EP, posterior endoderm. **b.** Samples for Geo-seq: embryonic stages, biological replicates and the germ layer cells. Stages: E6.5-E6.75, early-streak stage; E7.0-E7.25, mid-to late-streak stage; E7.5, late-streak to no-bud stage. **c.** The strategy of sampling of cell populations in the epiblast/ectoderm and endoderm in E6.5 to E7.5 embryos and mesoderm in E7.0 to E7.5 embryos (areas of sampling shown in histology images). Samples were designated in ascending order of serial sections (1 = the most distal section) and the regions in the section (R, reference section; S, sample section). Embryonic axes: anterior – posterior, A ↔ P; proximal – distal, P ↔ D; left – right, L ↔ R. Scale bar, 50 μm. **d.** Corn plots showing the spatio-temporal pattern of expression of *T* and *Otx2*. *T* expression domain marks the length of the primitive streak for staging the development of the embryo; *Otx2* is a representative ectoderm marker. **e.** Box plot showing the number of detected genes (FPKM > 0) in samples of E6.5- E7.5 embryos (1 biological replicate for E6.5, E6.75 and E7.25 embryos; 2 biological replicates for E7.0 and E7.5 embryos). The center line marks the median and box edges represent 25th and 75th percentiles. The median of genes detected in reference embryo per stage is 11,033 (E6.5), 10,150 (E6.75), 12,340 (E7.0), 12,549 (E7.25) and 14,246 (E7.5). **f.** Gene expression density plot of Geo-seq data of samples of E6.5-E7.5 embryos. The X-axis of the density plots is harmonized to the same scale. **g.** Saturation analysis for reads for samples from each development stage. Different numbers of reads were selected, and the number of detected genes was plotted.

**Extended Data Figure 2.**
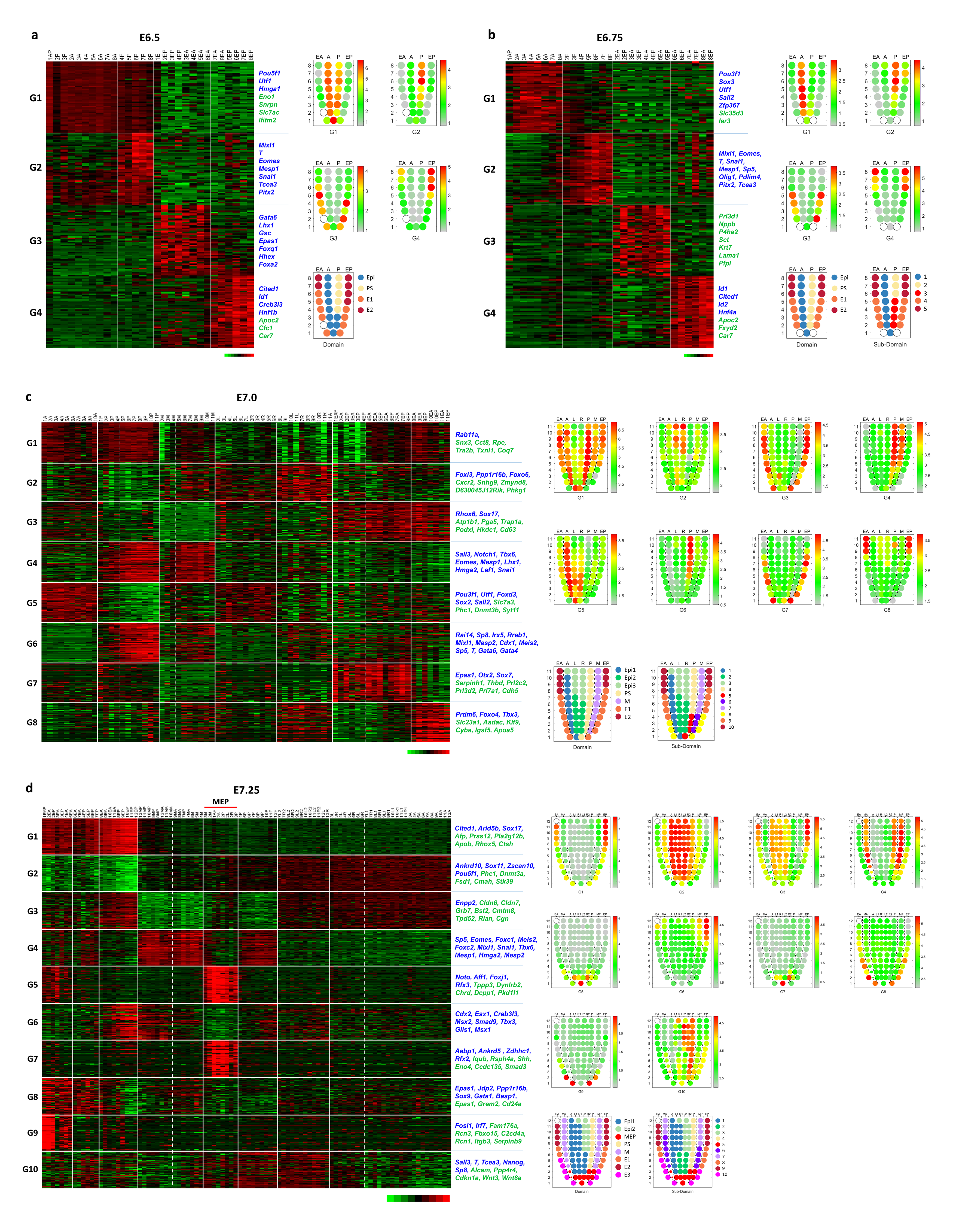
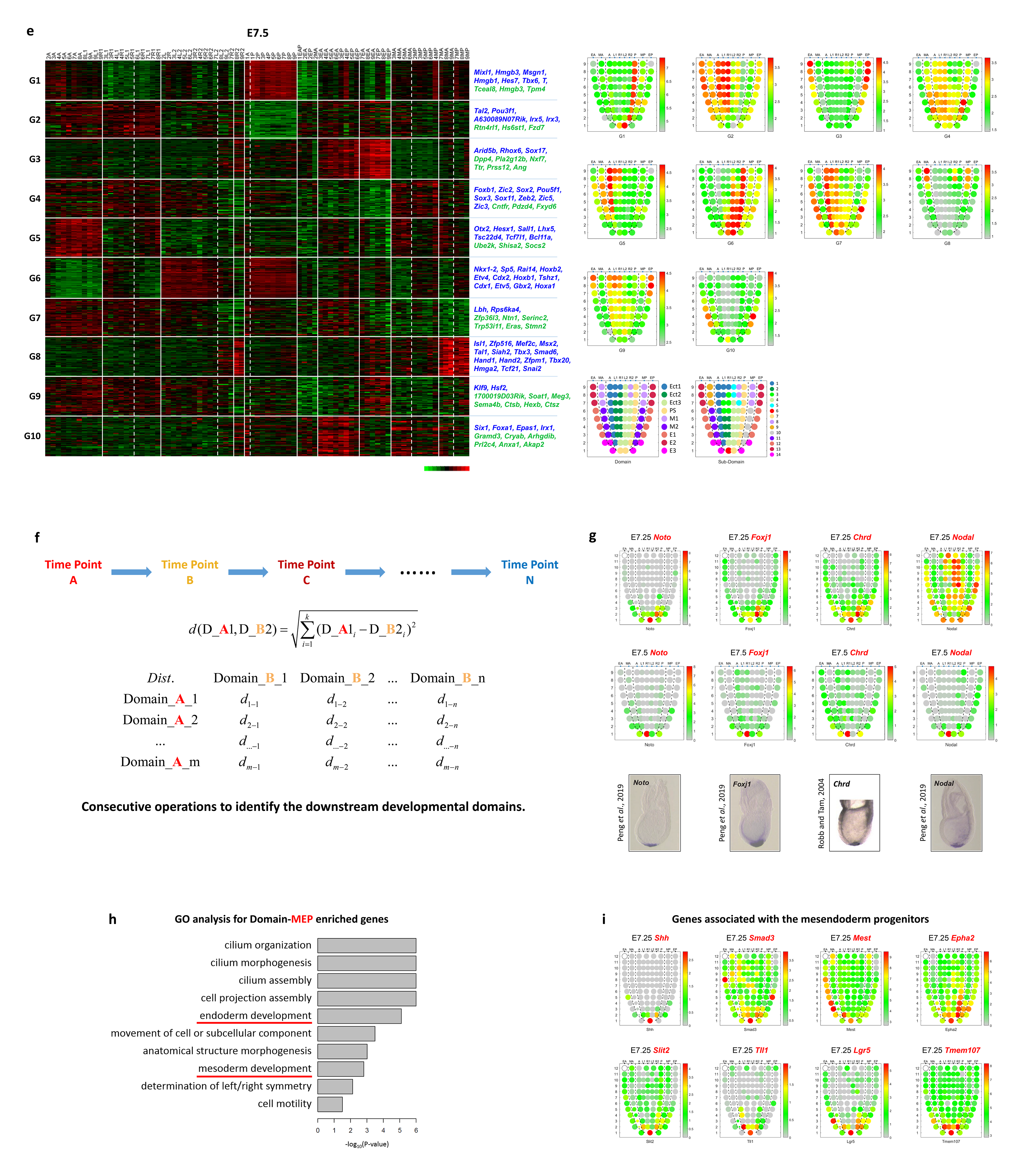
The spatial domains in the germ layers of E6.5-E7.5 embryos delimited by the expression profile of zipcode genes. **a-e.** Heat maps and corn plots of zipcode groups in E6.5 (**a**), E6.75 (**b**), E7.0 (**c**), E7.25 (**d**) and E7.5 (**e**) embryos. Transcription factors (blue marked) and top DEGs (green marked) of each gene group were listed on the right side of heat map. G, group. **f.** Population Tracing algorithm. The Euclidean distance of any two domains in embryos of successive stages were computed, following by delineating the mutual nearest neighbors of each domain from one stage against the domains of embryo at the successive stage. D_A1 denotes domain 1 at time point A, D_B2 denotes domain 2 at time point B. **g.** Expression pattern of node-related genes in E7.25 and E7.5 embryo. The spatial transcriptome (corn plot) data of *Noto*, *Foxj1*, *Chrd* and *Nodal* were verified with reference to whole mount in situ hybridization (WISH) data. WISH images (*Noto*, *Foxj1* and *Nodal*) were obtained from our previous study (Peng *et al*., 2019), WISH image of *Chrd* was obtained from a previous study (Robb and Tam, 2004). **h.** Functional ontology of enriched genes associated with the mesendoderm progenitors (MEP). **i.** Corn plots showing representative genes associated with the mesendoderm progenitors

**Extended Data Figure 3.**
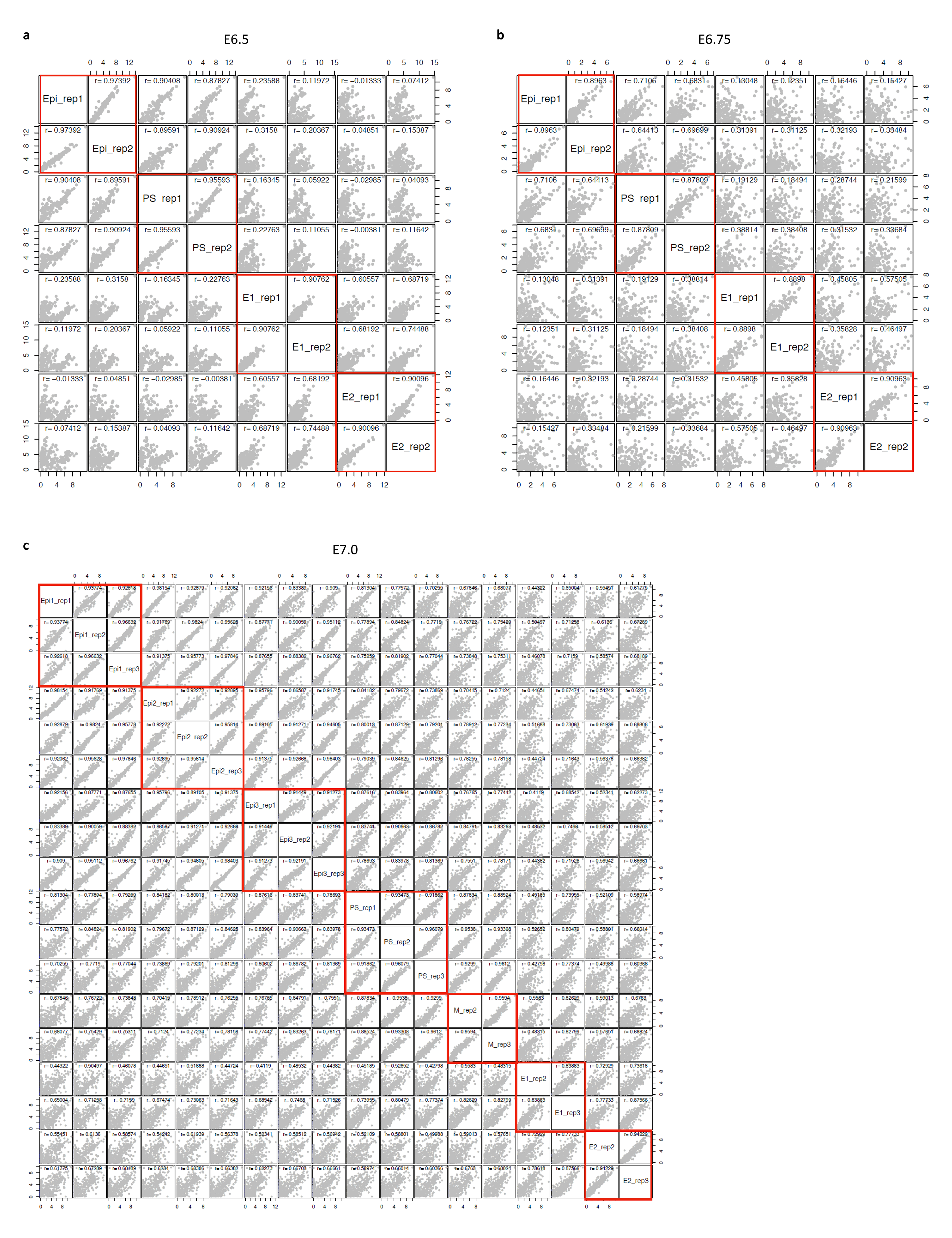
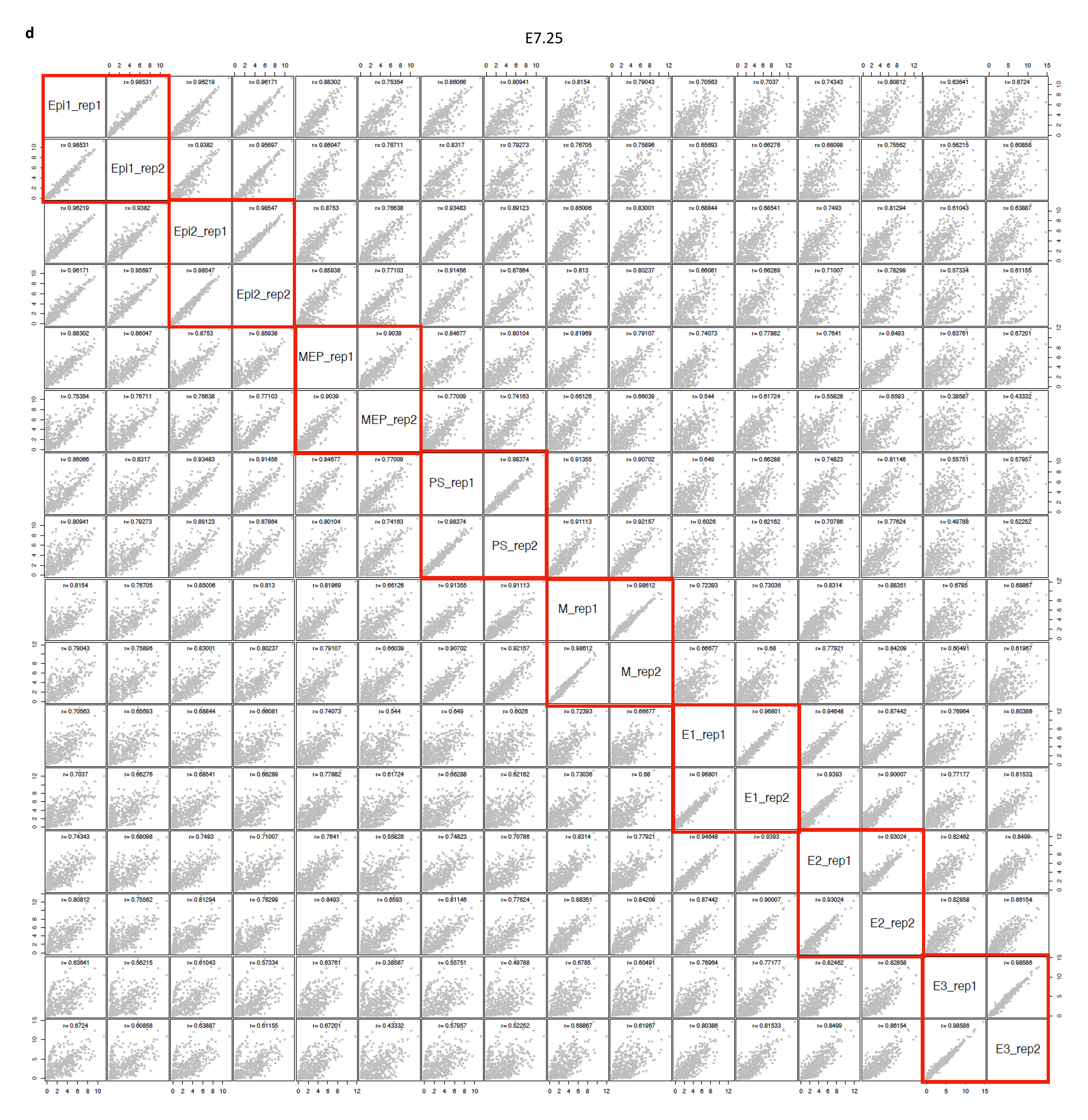
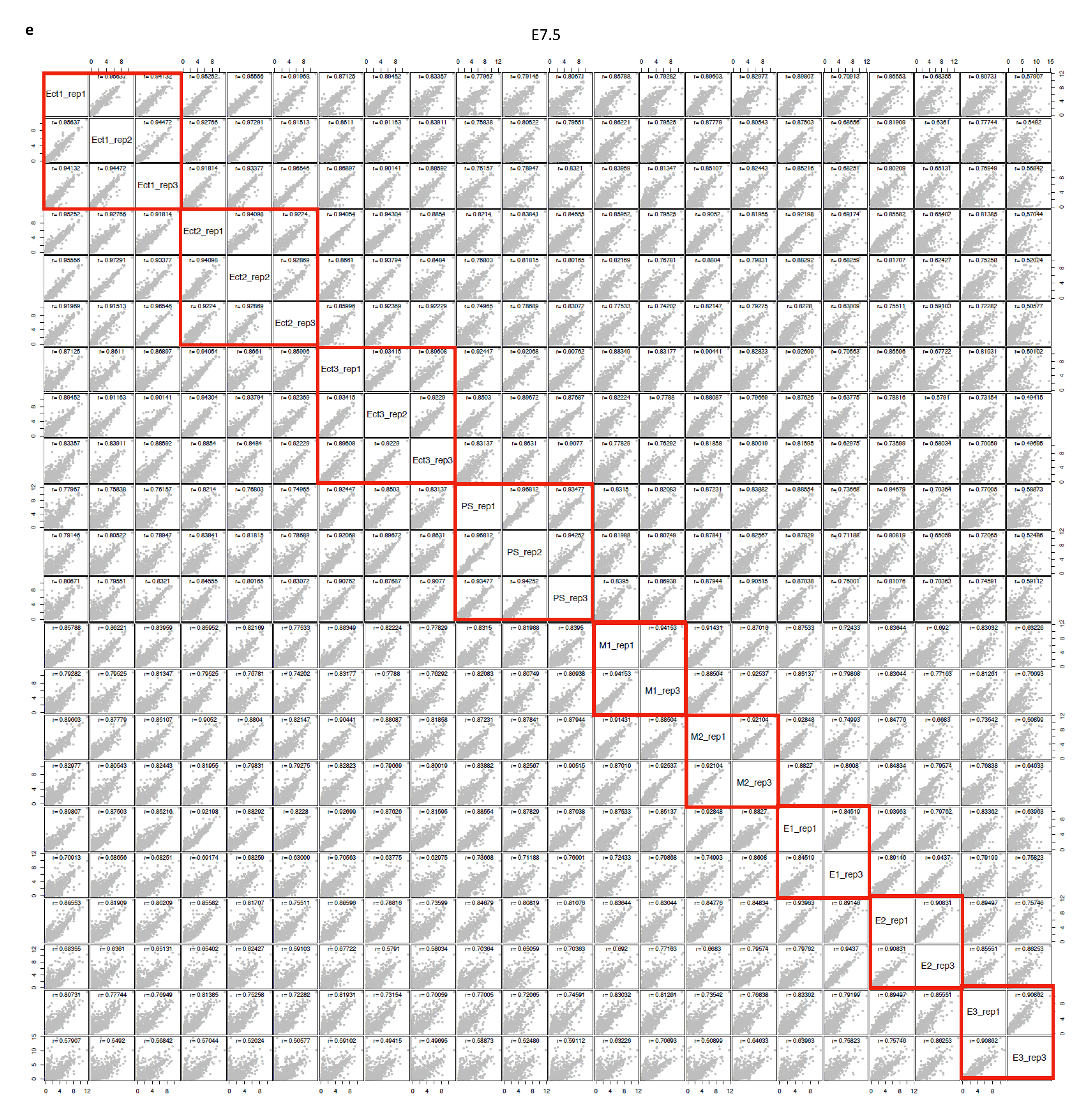
Inter-embryo correlation of spatial transcriptome data. **a-e.** The Pearson Correlation Coefficient (PCC) of spatial domains between biological replicates at E6.5 (**a**), E6.75 (**b**), E7.0 (**c**), E7.25 (**d**) and E7.5 (**e**). For E7.0 replicate 2 and E7.5 replicate 2, only epiblast/ectoderm domains were assessed.

**Extended Data Figure 4.**
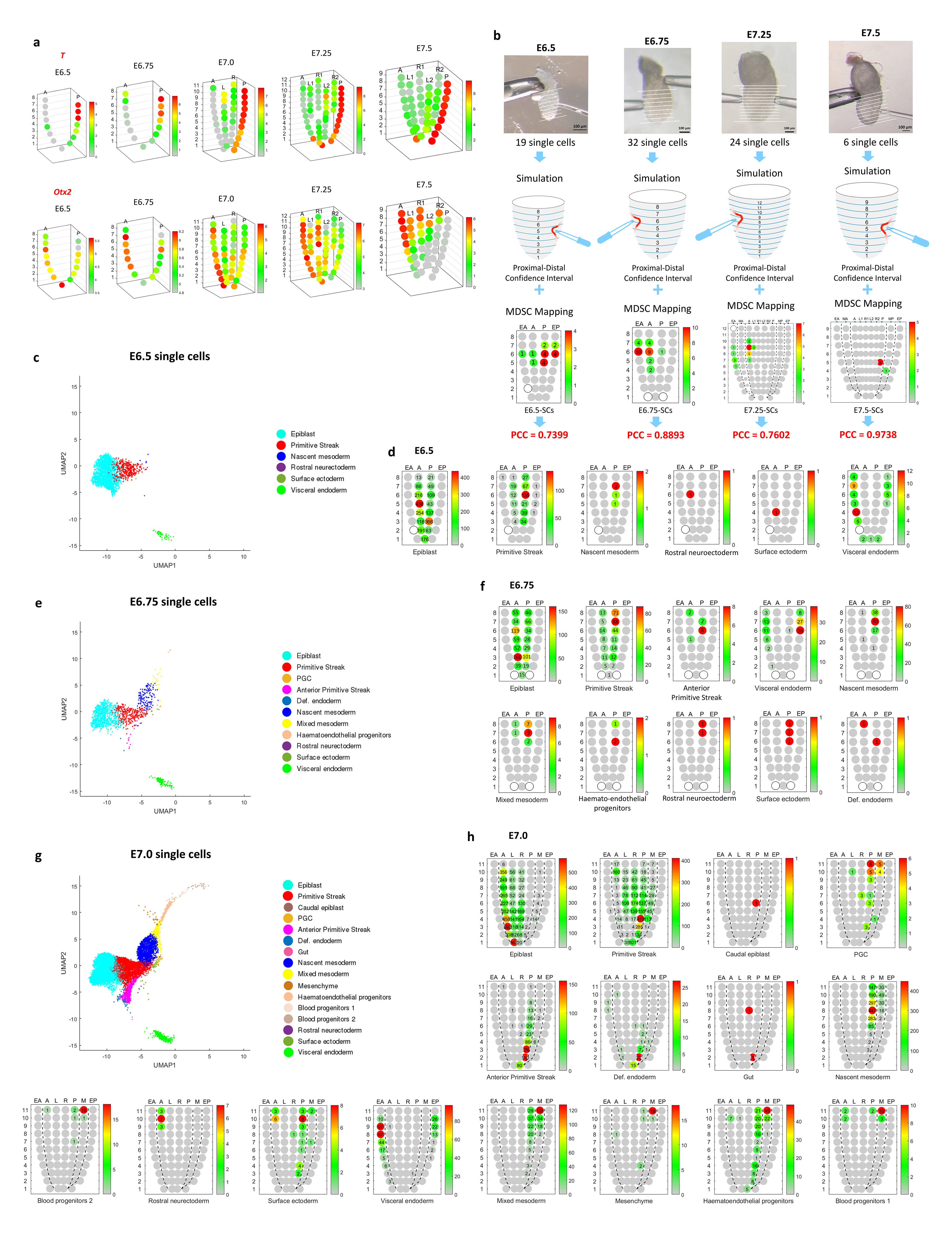
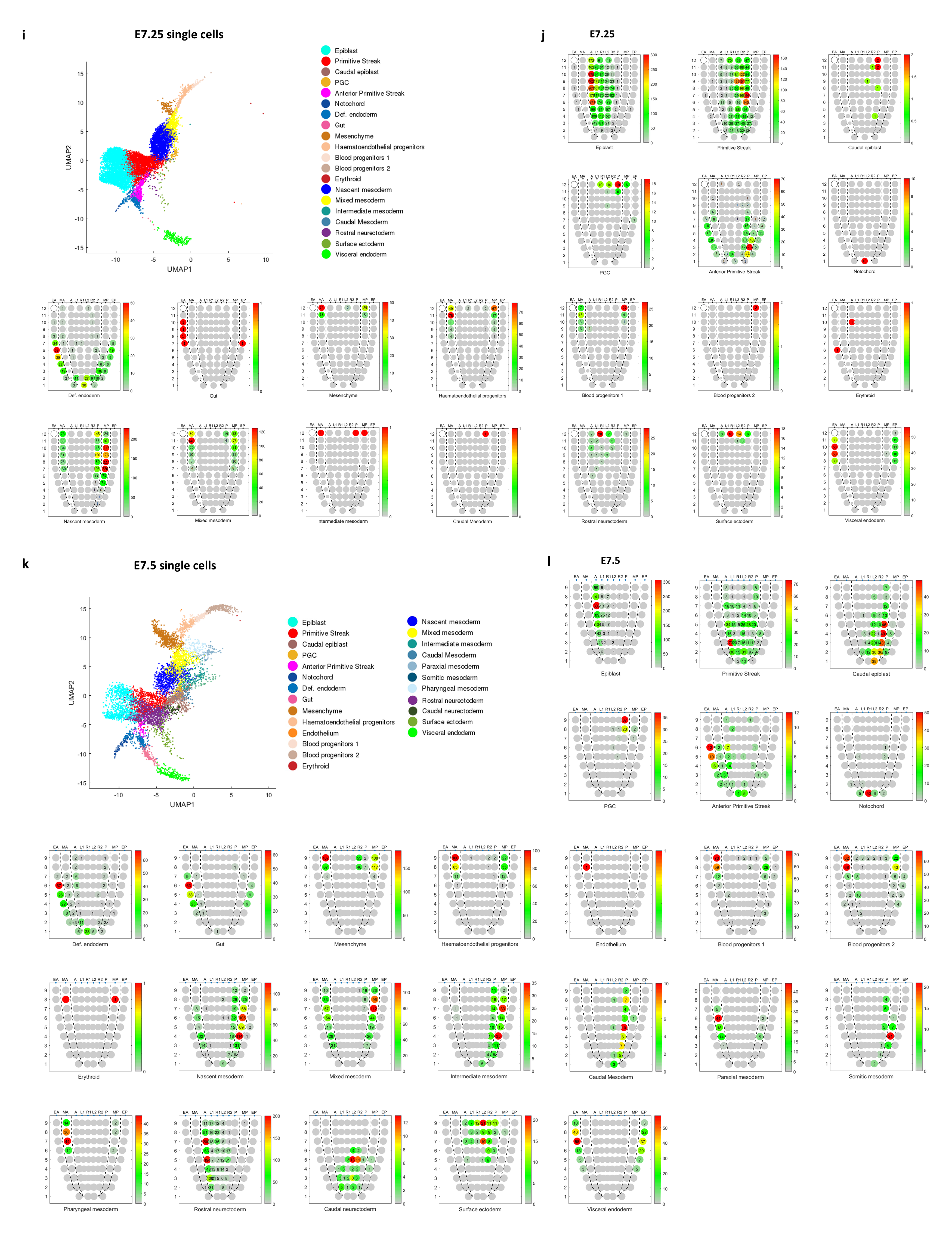
3D model for displaying transcriptome data and the verification of MDSC Mapping results. **a.** 3D corn plots showing the spatio-temporal pattern of expression of *T* and *Otx2* in epiblast/ectoderm layer of E6.5-E7.5 embryos. **b.** Verification of the results of MDSC Mapping of single cells isolated from known positions of E6.5, E6.75, E7.25 and E7.5 embryos. The number on each corn indicates the number of cells mapped to the specific position in the germ layers. PCC values and confidence intervals shown in the simulation. **c-l.** Uniform manifold approximation and projection (UMAP) plots (**c, e, g, i, k**) showing the data structure of single cells identified in the ‘Gastrulation Atlas’ and MDSC Mapping results (**d, f, h, j, l**,) for E6.5 (**c, d**), E6.75 (**e, f**), E7.0 (**g, h**), E7.25 (**i, j**) and E7.5 (**k, l**) embryos. Cell types are annotated (legend of UMAP) and the spatial distribution of each annotated cell types is displayed in corn plots, with the number of cells mapped to specific Geo-seq position (number in the corn) shown.

**Extended Data Figure 5.**
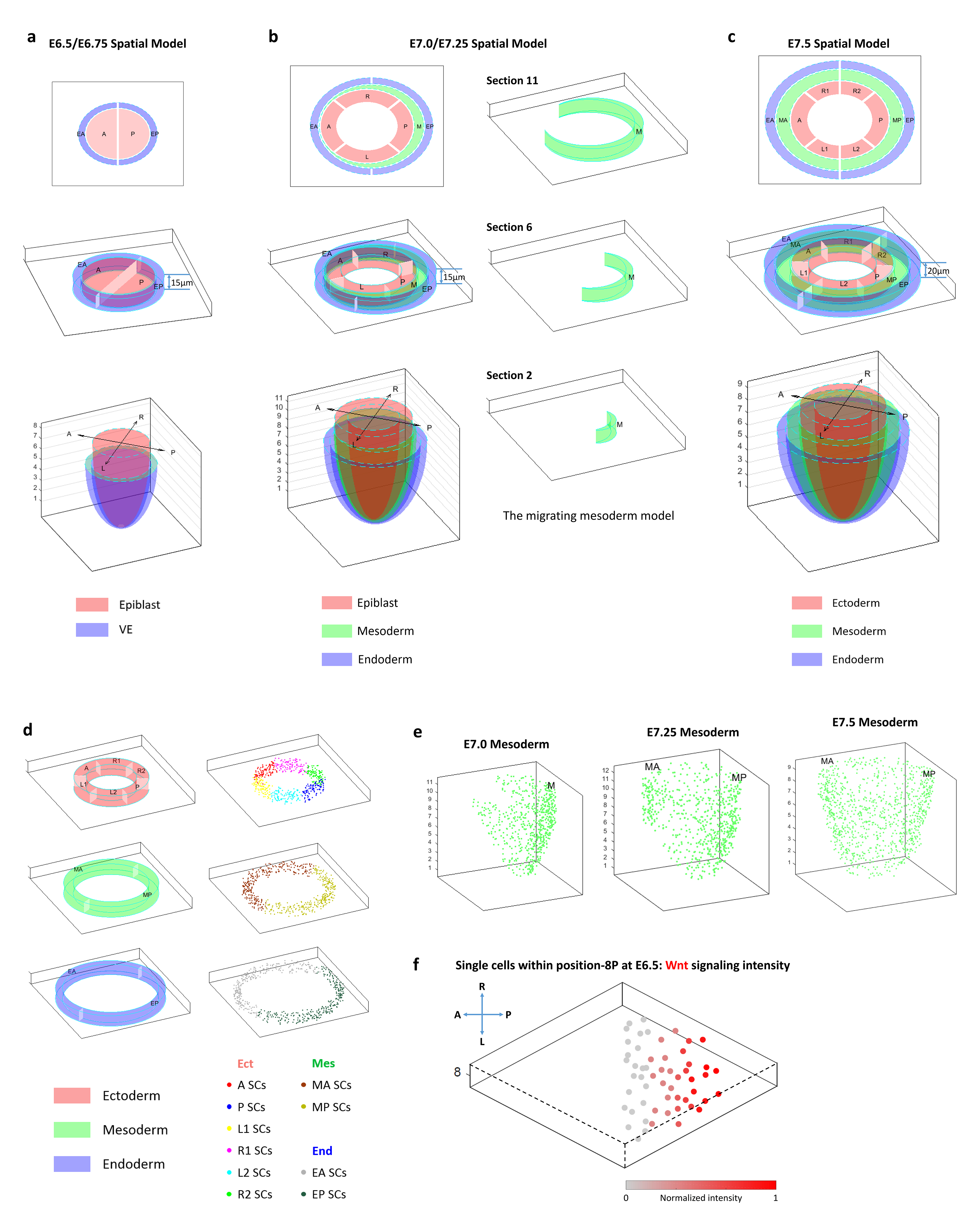
3D modeling for single-cell resolution embryo map. **a.** The 3D model of E6.5 and E6.75 embryos. Based on Geo-seq sampling strategy, semicircular pattern for the epiblast and Annulus Model for the endoderm were devised to display the spatial distribution of single cells. Section thickness, 15 μm. **b.** The Annulus Model of the epiblast and endoderm of E7.0 and E7.25 embryos. At E7.0-E7.25, the Migration Model mirrors the mesoderm layer spanning from posterior to anterior of the embryo. Section thickness, 15 μm. **c.** The Annulus Model of E7.5 embryo. Concentric annuli represent the ectoderm, mesoderm and endoderm from the inside outward. Section thickness, 20 μm. **d.** The display of single cells in the three germ layers of E7.5 embryo. Single cells that mapped to a Geo-seq position were distributed uniformly across the corresponding interior space of each domain in the annulus section. **e.** The display of single cells in the Migration Model of the mesoderm of E7.0-E7.5 embryo. **f.** Bubble Sort algorithm re-ordered the distribution pattern of single cells in E6.5 position-8P by graded WNT activity in the anterior-posterior axis. The color legend indicates the normalized expression level determined by averaged transcript counts.

**Extended Data Figure 6.**
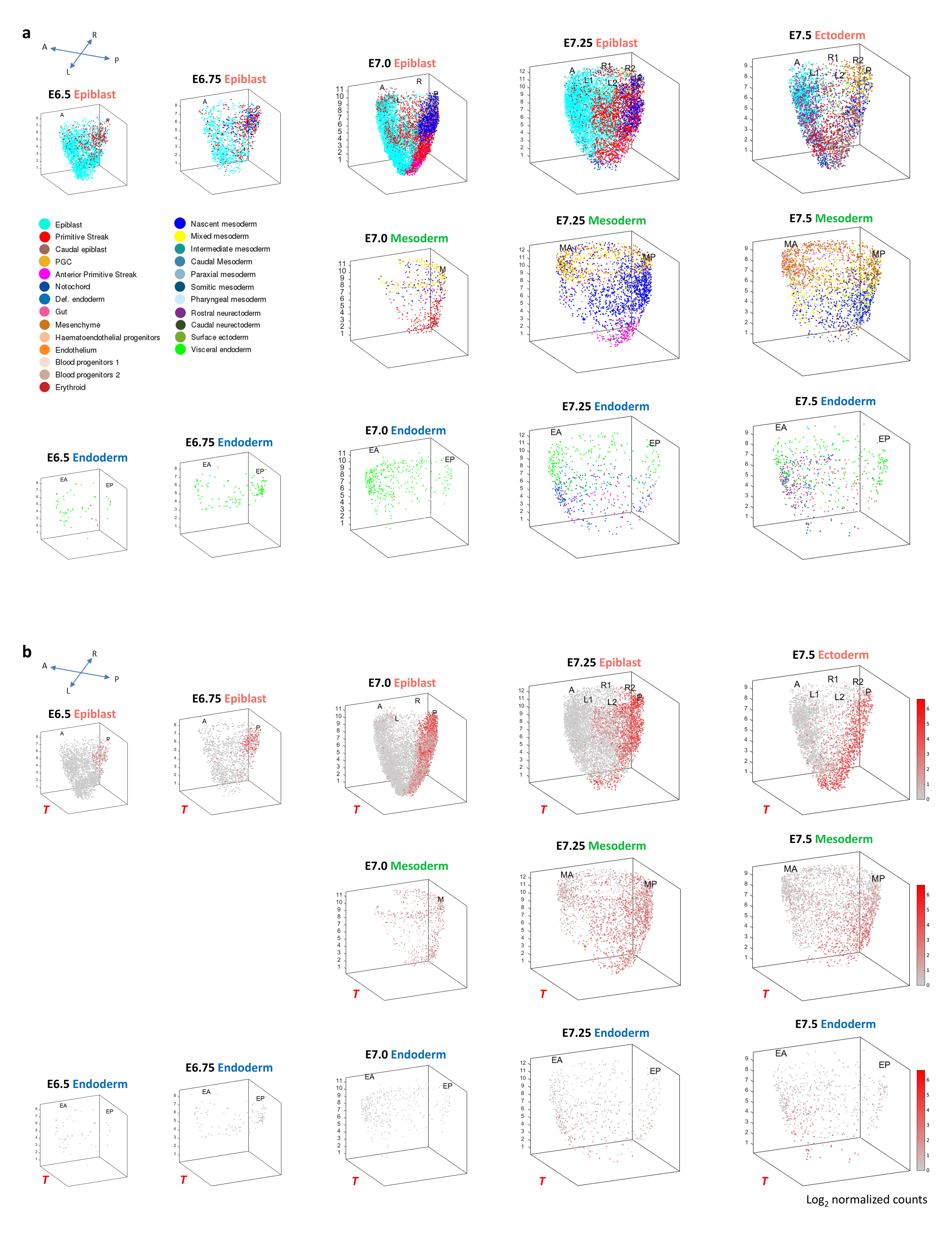
The spatio-temporal distribution of single cells in E6.5- E7.5 mouse embryos. **a.** The spatio-temporal distribution of all the single cells identified in the ‘Gastrulation Atlas’ in the epiblast/ectoderm, mesoderm and endoderm of E6.5- E7.5 mouse embryos. **b.** The spatio-temporal distribution of *T*-expressing cells in E6.5-E7.5 embryos. The color legend indicates the level of expression determined by the transcript counts.

**Extended Data Figure 7.**
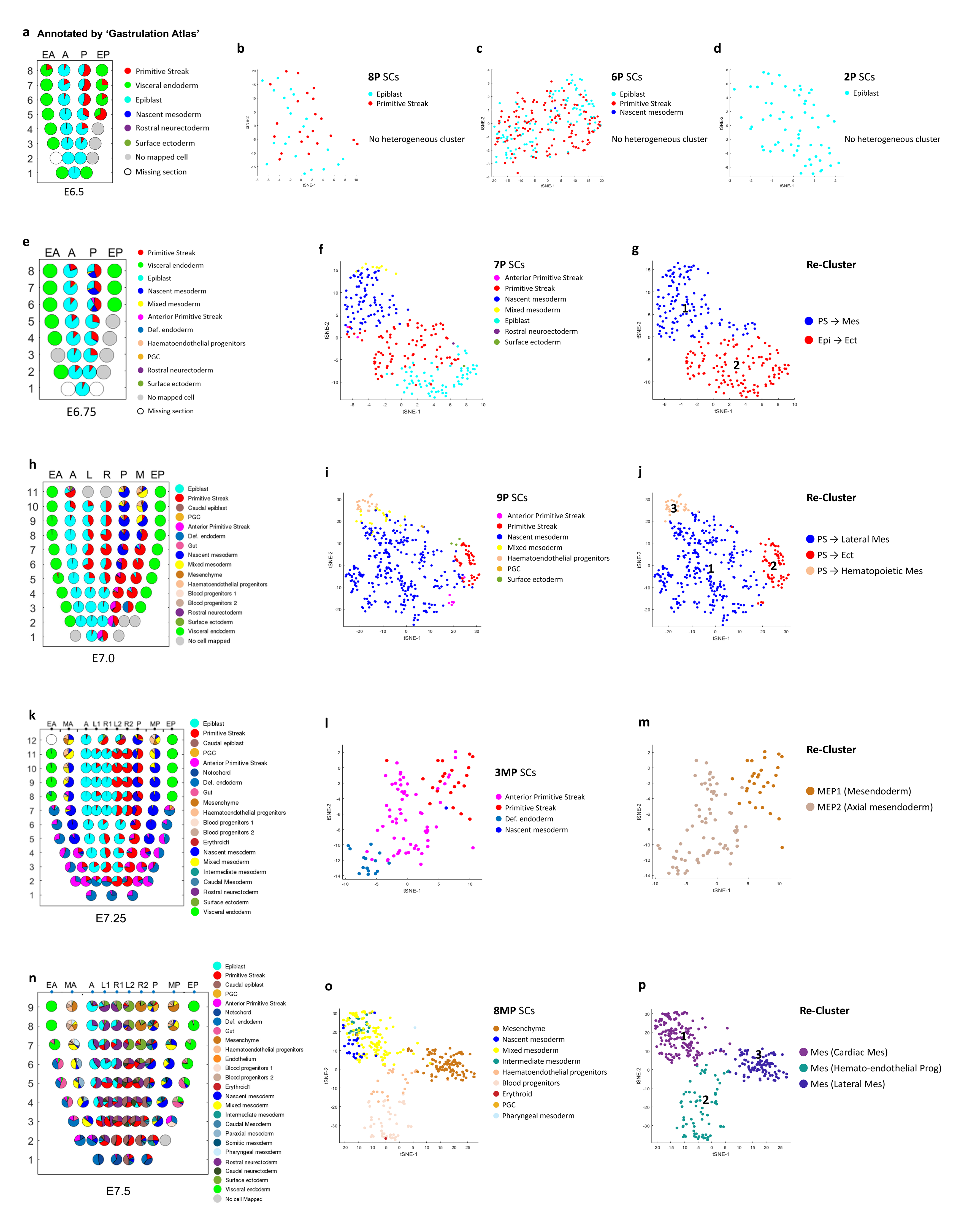
Analysis of heterogeneity of single cell population. **a-p.** Heterogeneity Map of single cell types annotated by ‘Gastrulation Atlas’ displayed as pie charts in corn plots (**a, e, h, k, n**) and *t*-SNE plots of single cells (**b, c, d, f, g, i, j, l, m, o, p**) showing examples of heterogeneity of cell types (with revised annotation according to **Extended Data** Fig. 8) in specified Geo-seq positions of E6.5 (**a-d**), E6.75 (**e-g**), E7.0 (**h-j**), E7.25 (**k-m**) and E7.5 (**n-p**) embryos.

**Extended Data Figure 8.**
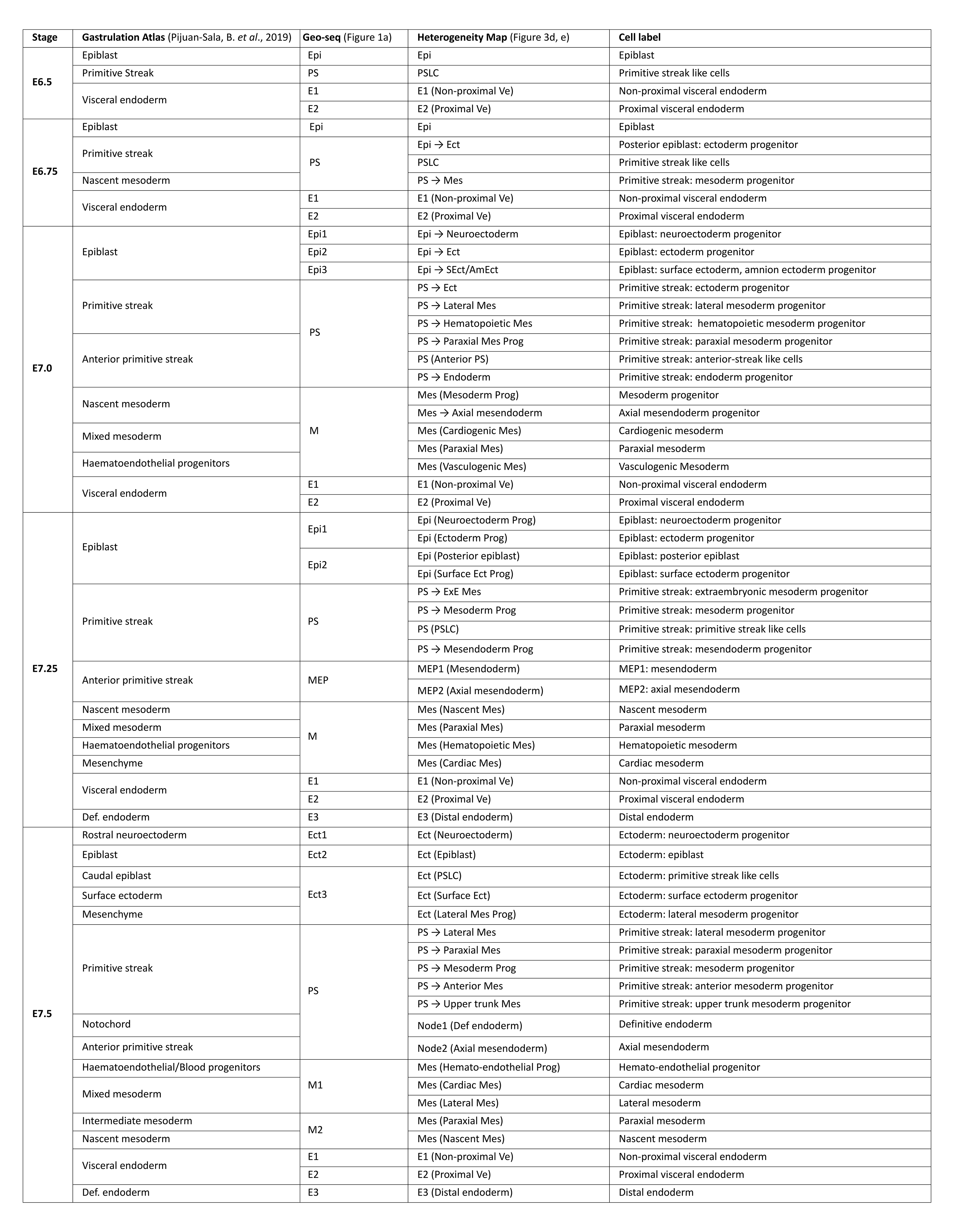
Re-annotation of cell types. Revised annotation of single cell types mapped to the Geo-seq position in the epiblast/ectoderm, primitive streak, mesoderm and endoderm. Annotation of cell types in ‘Gastrulation Atlas’ is shown for comparison. The nomenclature ‘X→Y’ and ‘X(Y)’ represent different cell states. X represents the germ layer information of cell population. ‘X→Y’ indicates these cells are representing a transitional cell state from X to Y. And ‘X(Y)’ represents the precursor of a specified cell type (Y) in the germ layer X.

**Extended Data Figure 9.**
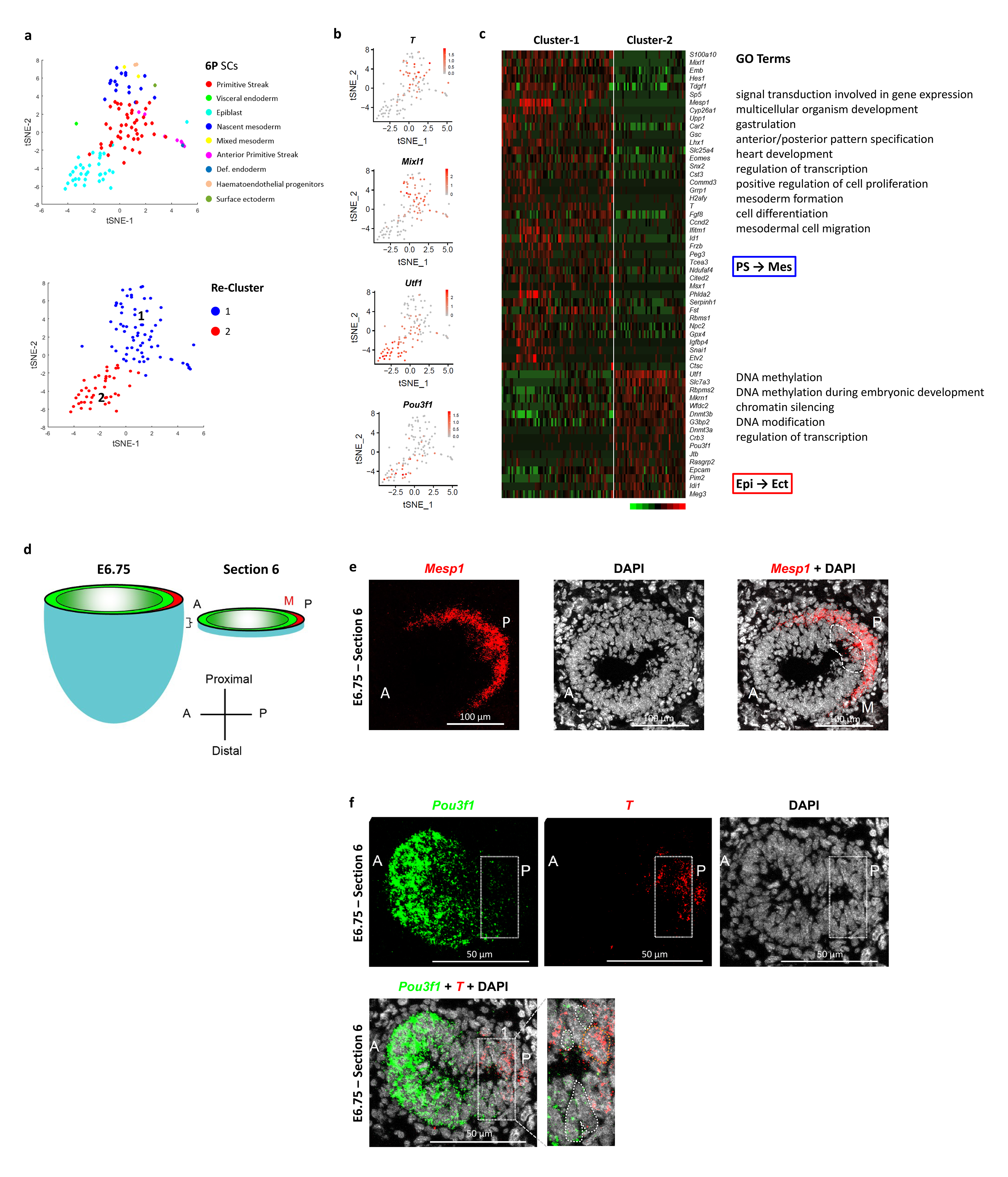
Heterogeneity analysis and validation of marker genes. **a.** *t*-SNE plot showing the single cells mapped to position-6P at E6.75 (top panel). Re-clustering revealed the presence of two distinct clusters (bottom panel). **b.** *t*-SNE plot showing the expression pattern of *T* and *Mixl1* for cluster-1, and *Utf1* and *Pou3f1* for cluster-2. **c.** Heat map showing the differentially expressed genes of the two cell clusters (p < 0.01, fold change > 1.5). The enriched gene ontology (GO) terms (p < 0.05) provided additional information for annotating Cluster-1 as ‘PS→Mes’ and Cluster-2 as ‘PS →Ect’. **d.** RNAscope analysis for E6.75 embryo. **e.** RNAscope analysis validated the expression of *Mesp1* in the posterior epiblast of E6.75 embryo. **f.** RNAscope analysis validated the co-localization of *T* and *Pou3f1* at the posterior epiblast of E6.75 embryo.

**Extended Data Figure 10.**
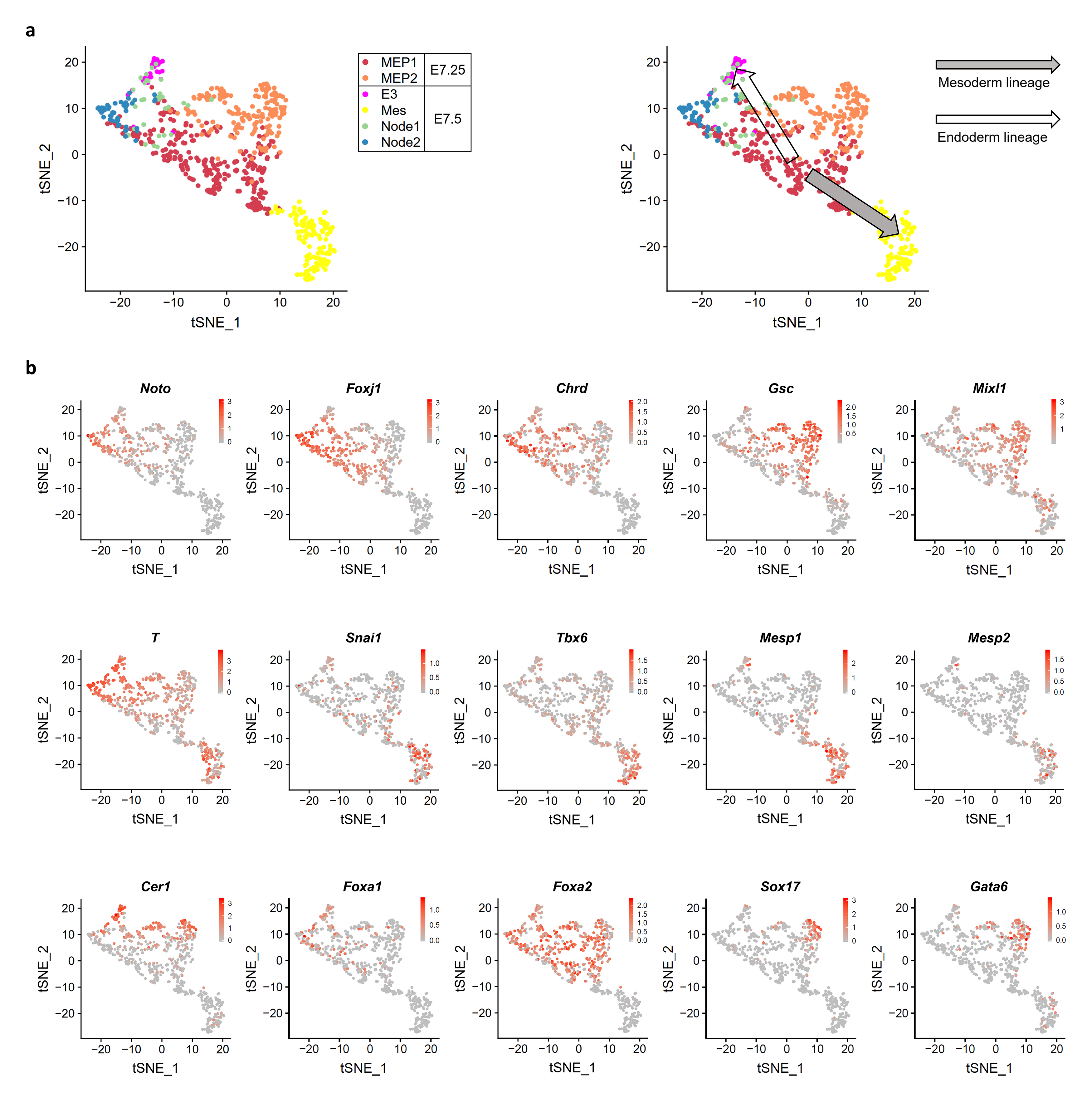
Imputation of molecular trajectories. **a.** *t*-SNE plots showing the E7.25 putative MEPs and the inferred E7.5 derivatives. Left panel: annotated cell types, right panel: developmental trajectory. **b.** *t*-SNE plots showing the expression pattern of marker genes of node (top panels), mesoderm (middle panels) and endoderm (bottom panels). The color legend indicates the level of expression determined by transcript counts.

**Extended Data Figure 11.**
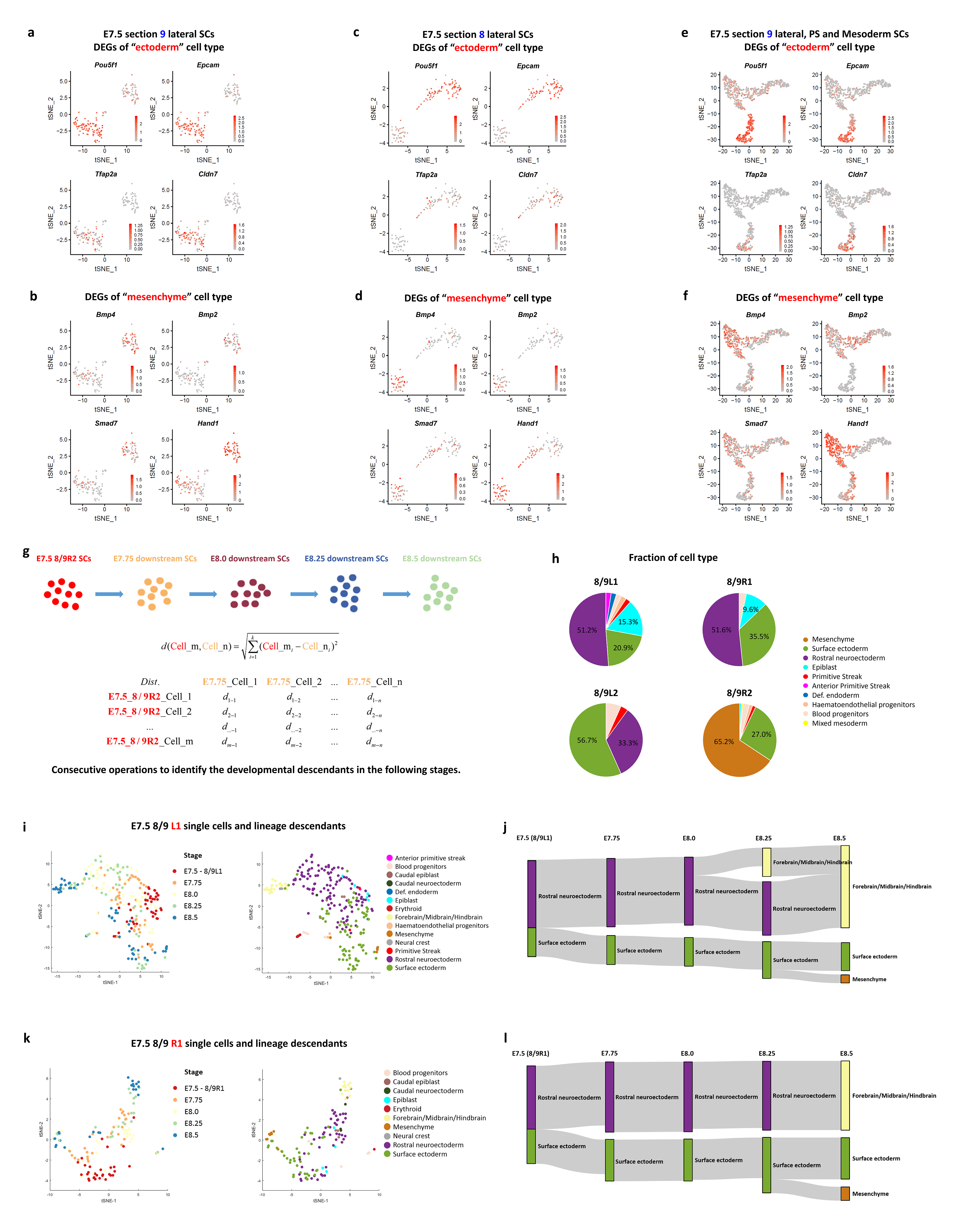
Imputation of molecular trajectories. **a-f.** *t*-SNE plots showing the differentially expressed genes of “ectoderm” cell type (**a, c, e**) and “mesenchyme” cell type (**b, d, f**) in position-8R2/9R2 (corresponding to Fig. 4a**-f**). Panel **a**-**d** showed the single cells of proximal-lateral ectoderm positions in section-9 (**a**, **b**) and −8 (**c**, **d**). Panel **e**, **f** showed the single cells of proximal-lateral ectoderm, primitive streak and mesoderm positions in section-9. **g.** Schematics of Population Tracing algorithm for single cells of E7.5-E8.5 embryos. **h.** Pie charts showing the fraction of cell types at position-8L1/9L1, 8R1/9R1, 8L2/9L2 and 8R2/9R2. **i-l.** *t*-SNE plots (**i, k**) and the molecular trajectories (**j, l**) of single cells (imputed using the Population Tracing algorithm) at position-8L1/9L1 (**i, j**) and position-8R1/9R1 (**k**, **l**) of E7.5 embryo and cells in E7.75-E8.5 embryos. Developmental timepoints (stage) and cell types (see legend of panel **i**, **k**) are indicated in the *t*-SNE plots. Cell types in E7.75-E8.5 embryos are annotated according to the ‘Gastrulation Atlas’.

**Extended Data Figure 12.**
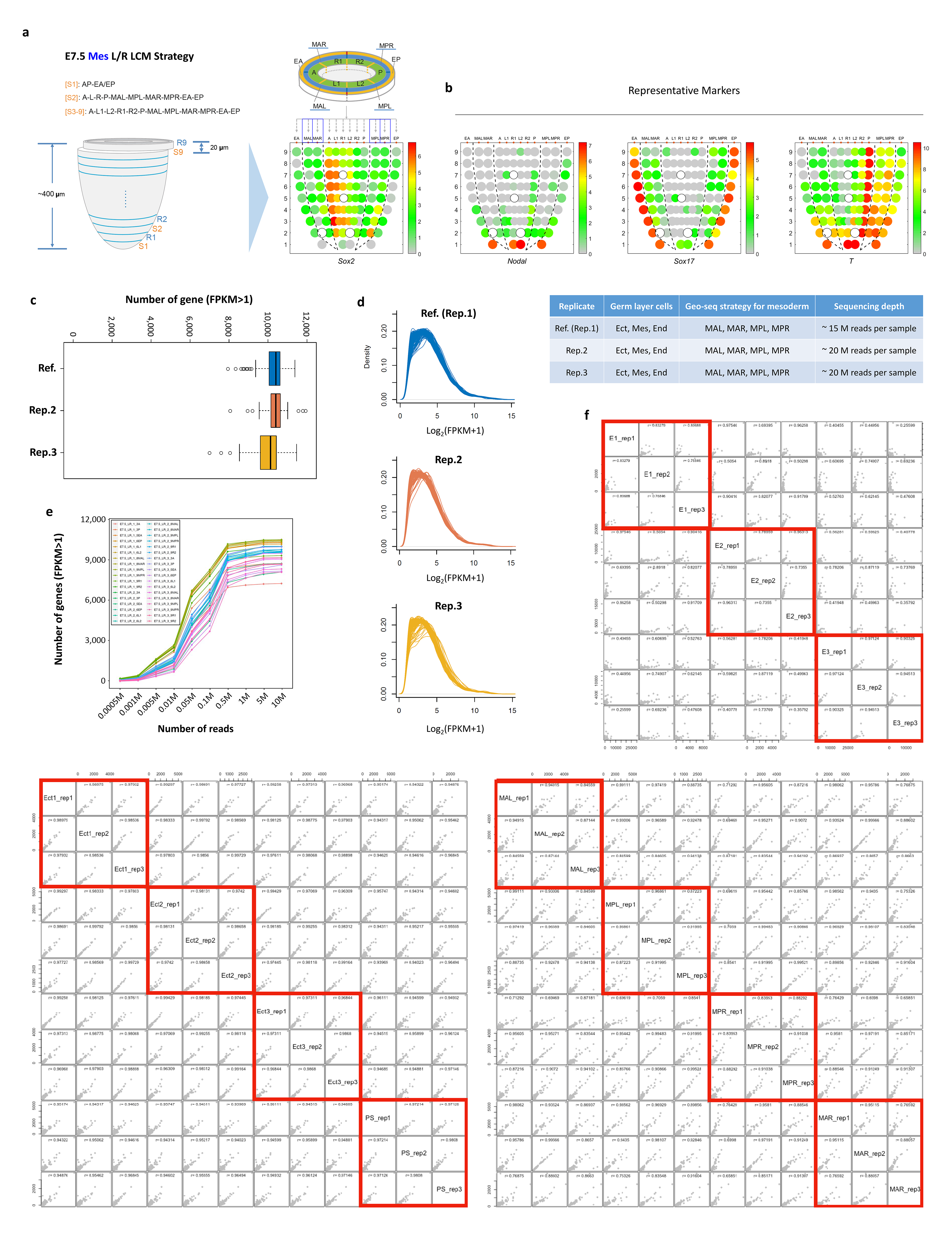
Refined Geo-seq analysis of the mesoderm cell layer of the E7.5 embryo. **a.** The strategy of sampling cell populations in the E7.5 embryo and the presentation of the spatial pattern of gene expression in the corn plot. MAL, anterior left mesoderm; MAR, anterior right mesoderm; MPL, posterior left mesoderm; MPR, posterior right mesoderm. **b.** Corn plots showing the spatial pattern of expression of representative marker genes: *Nodal*, *Sox17*, *T*. Hollow circles indicate missing samples. Table: Geo-seq strategy and sequencing depth for biological replicates. **c.** Box plot showing the number of detected genes (FPKM > 1) in the 3 biological replicates. The center line marks the median and box edges represent 25th and 75th percentiles. The median genes detected per replicate is 10,358 (Ref.), 10,389 (Rep. 2) and 10,042 (Rep. 3). **d.** Gene expression density plot of Geo-seq data of the 3 biological replicates. **e.** Saturation analysis. Different numbers of reads were selected, and the number of detected genes was plotted. **f.** Inter-embryo correlation of spatial transcriptome data. The Pearson Correlation Coefficient (PCC) of spatial domains between biological replicates were calculated and showed high inter-embryo consistencies.

**Extended Data Figure 13.**
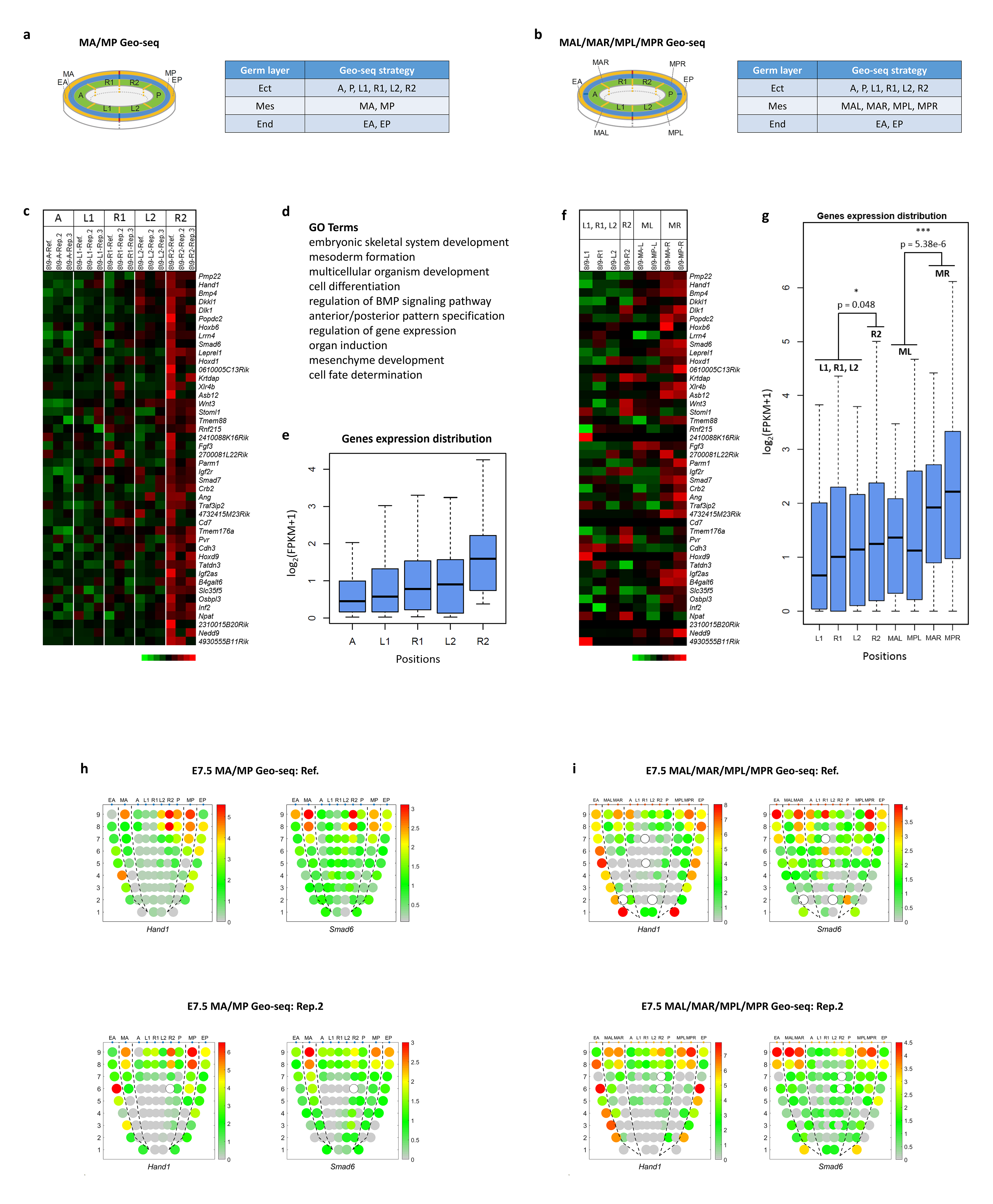
The right-side DEGs of the proximal-lateral ectoderm showed more significant differences in the proximal-lateral mesoderm. **a.** The Geo-seq strategy of sampling cells from MA/MP populations in the E7.5 embryo. MA, anterior mesoderm; MP, posterior mesoderm. **b.** The Geo-seq strategy of sampling cells from MAL/MAR//MPL/MPR populations in the E7.5 embryo. MAL, anterior left mesoderm; MAR, anterior right mesoderm; MPL, posterior left mesoderm; MPR, posterior right mesoderm. **c.** Heatmap showing the differentially expressed genes (DEGs) in 8/9R2 regions compared with other lateral regions (8/9L1, 8/9L2, and 8/9R1) and anterior (A) region in the 3 Geo-seq replicates (Original MA/MP Geo-seq strategy), p < 0.1, fold change > 1.5. **d.** The enriched gene ontology (GO) terms for the 8/9R2 specific DEGs. **e.** Box plot showing the distribution of expression level (Log2 normalized FPKM) of 8/9R2 specific DEGs in different ectoderm regions. **f.** Heatmap showing the expression pattern of 8/9R2 specific DEGs (identified in **c**) in different regions of the MAL/MAR/MPL/MPR Geo-seq embryo. **g.** Box plot showing the distribution of expression level (Log2 normalized FPKM) of 8/9R2 specific DEGs in different regions of the MAL/MAR/MPL/MPR Geo-seq embryo. T-test p-values were labelled on the box plot. *, p < 0.05; **, p < 0.01; ***, p < 0.001. **h.** Corn plots showing the spatial expression pattern of *Hand1* and *Smad6* in the MA/MP Geo-seq embryos. Both reference and replicate embryos show L-R difference in the proximal ectoderm. **i.** Corn plots showing the spatial expression pattern of *Hand1* and *Smad6* in the MAL/MAR/MPL/MPR Geo-seq embryos. Both reference and replicate embryos show more significant difference in the right-side proximal lateral mesoderm.

**Extended Data Figure 14.**
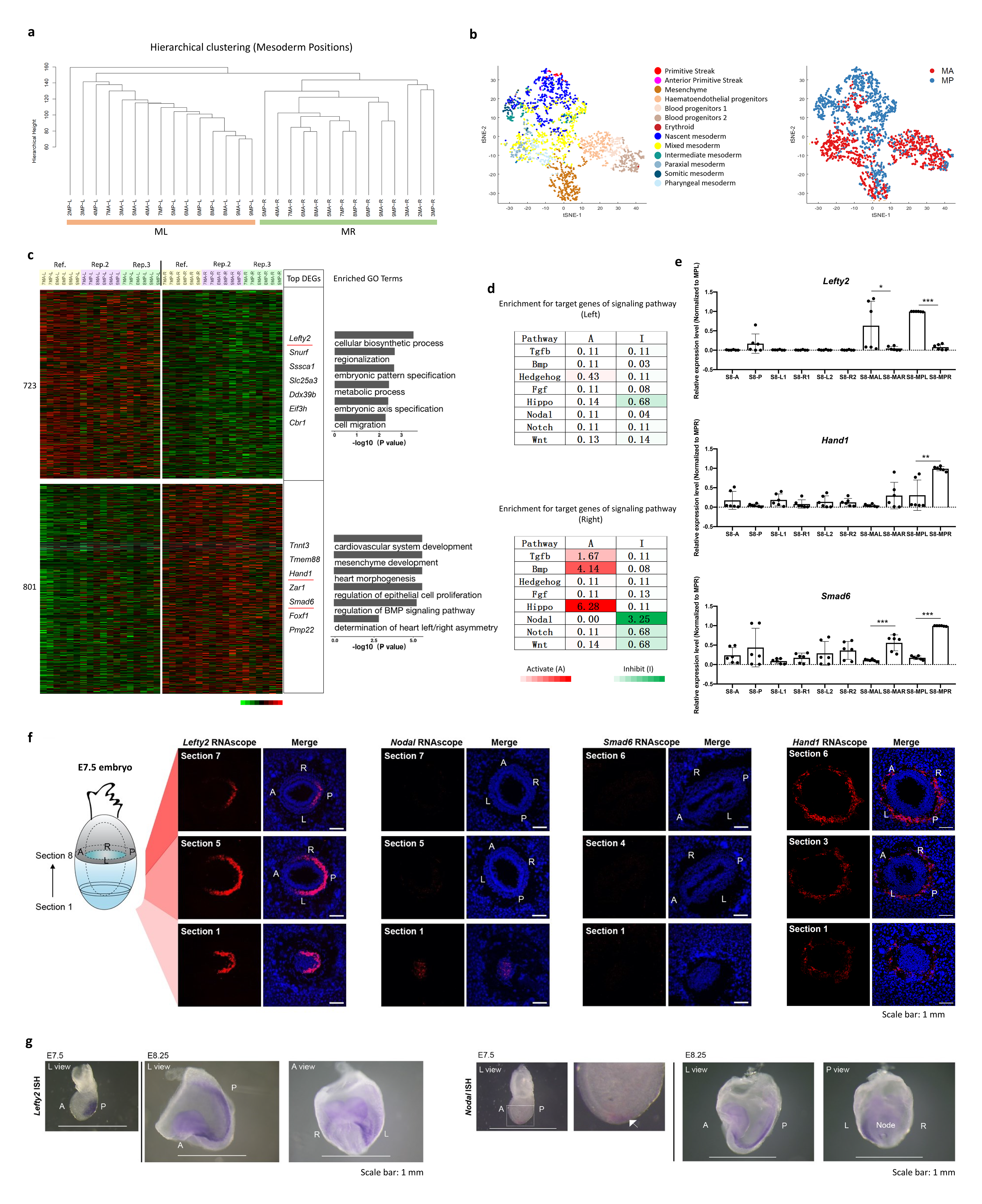
Validation for the left-right asymmetric genes. **a.** Hierarchical clustering showed the difference between the mesoderm tissues on contralateral sides of the E7.5 embryo. **b.** *t*-SNE plot showing the data structure of single cells that are allocated to mesoderm positions at E7.5. Cells are colored by both their cell-type annotation according to ‘Gastrulation Atlas’ (left) and anterior/posterior location based on MDSC Mapping results (right). **c.** Heat map showing the differentially expressed genes (DEGs) of the proximal-left mesoderm (n = 723) and proximal-right mesoderm (n = 801) regions across the 3 Geo-seq replicates. The top DEGs and the enriched gene ontology (GO) terms for each group were listed on the right (p < 0.01). The genes with red underline were assayed by real-time PCR in **e**. **d.** The enrichment for target/response genes of development-related signaling pathways in the proximal-left and proximal-right mesoderm. Signaling activity: red, activating (A); green, inhibitory (I). The significance of –log_10_(FDR) value in each cell was calculated by one-sided Fisher’s exact test followed by Benjamini-Hochberg correction. **e.** Real-time PCR showing significant left/right differences of *Lefty2*, *Hand1* and *Smad6* expression in the proximal mesoderm (section 8) of E7.5 embryo. *, p < 0.05; **, p < 0.01; ***, p < 0.001. **f.** RNAscope analyses showing the expression of *Lefty2, Nodal, Smad6* and *Hand1* in specified transverse sections (numbered) of E7.5 embryo. These RNAscope analyses serve as negative controls to the results in Fig. 5h. In these sections, there is no discernible difference between lateral sides, except for *Lefty2* in Section 5 and 7. A, anterior; P, posterior; L, left; R, right. **g.** Whole-mount in situ hybridization (WISH) showed the laterally asymmetric expression pattern of *Lefty2* (left panel) and *Nodal* (right panel, arrow) in E7.5 and E8.25 mouse embryos. A, anterior; P, posterior; L, left; R, right.

**Extended Data Fig. 15.**
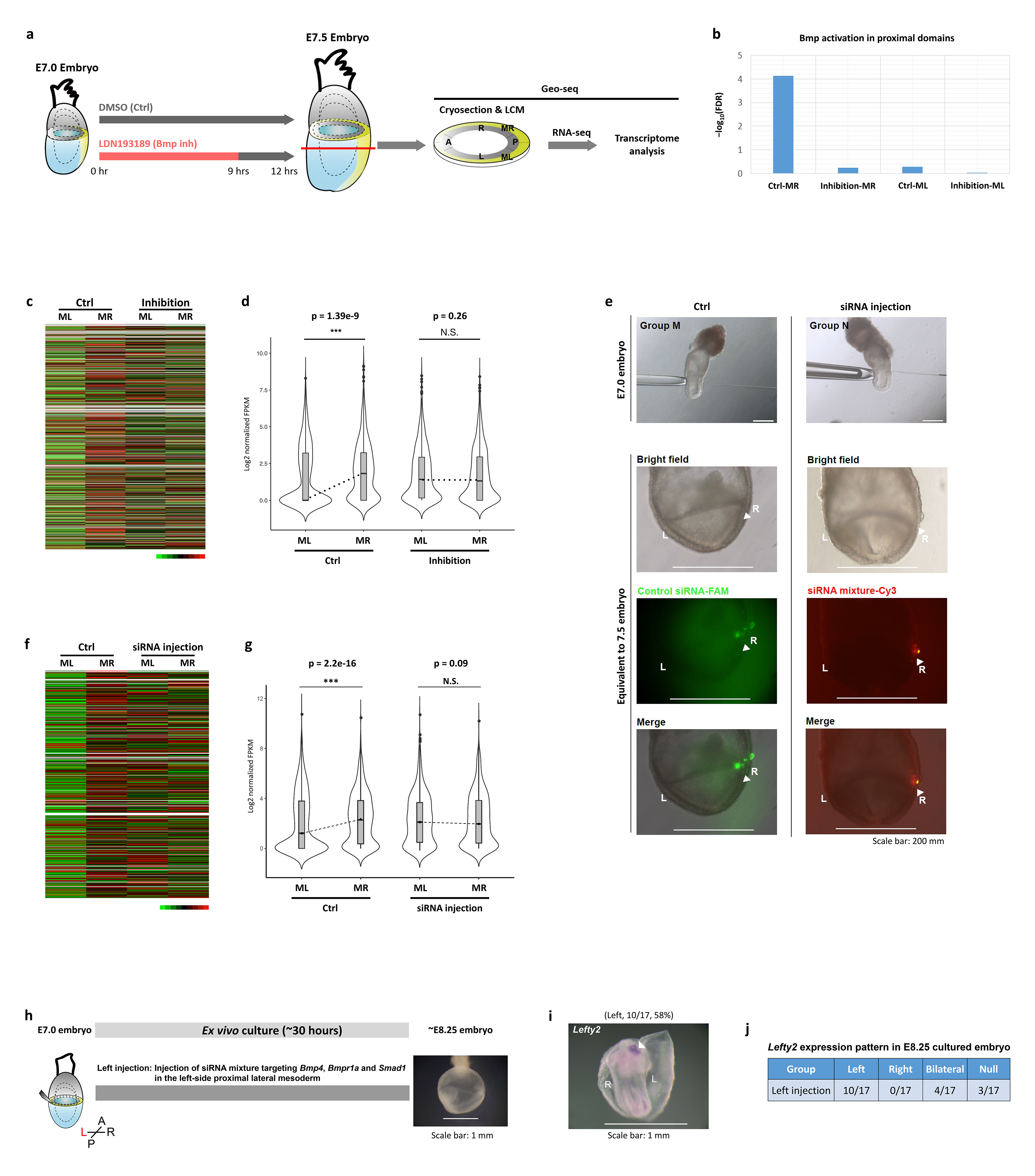
Geo-seq analyses for the *ex vivo* cultured embryos. **a.** Schematic showing the workflow of GEO-seq for *ex vivo* cultured embryos. **b.** The BMP signaling activity in the right-side lateral mesoderm region (MR) was curtailed by LDN193189 treatment. **c.** Heatmap showing the expression pattern of right-side specific genes of proximal lateral mesoderm (n = 801) in Control (Ctrl) and 9 hours BMP Inhibition embryos. A few right-side specific genes of proximal lateral mesoderm were not expressed in cultured embryos, these genes were then marked with white lines. **d.** Violin plot showing the expression level of right-side specific genes of proximal lateral mesoderm in control (Ctrl) and 9 hours BMP Inhibition embryos. The left-right differences were abrogated in the inhibition group. Rank sum test, ***, p < 0.001; N.S., no significant difference. **e.** Region-specific (the right-side proximal lateral mesoderm) perturbation of BMP signaling by multiple siRNA knockdowns. Top panel: The experiment snapshot of siRNAs microinjection. Bottom panel: Region-specific transfection of corresponding siRNAs can be observed at embryos of equivalent E7.5 stage. **f.** Heatmap showing the expression pattern of right-side specific genes of proximal lateral mesoderm (n = 801) in Control (Ctrl) and siRNA knockdown embryos. A few right-side specific genes of proximal lateral mesoderm were not expressed in cultured embryos, these genes were then marked with white lines. **g.** Violin plot showing the expression level of right-side specific genes of proximal lateral mesoderm in control (Ctrl) and siRNA knockdown embryos. The left-right differences were abrogated in the Bmp siRNA microinjection group. Rank sum test, ***, p < 0.001; N.S., no significant difference. **h.** Experimental strategy of siRNA knockdown in the left-side mesoderm and *ex vivo* culture. **i.** Whole-mount in situ hybridization of *Lefty2* of cultured embryo at equivalent E8.25 stage, showing *Lefty2* expression in the lateral mesoderm. **j.** Pattern of *Lefty2* expression in the cultured embryos at equivalent E8.25 stages.

**Extended Data Fig. 16.**
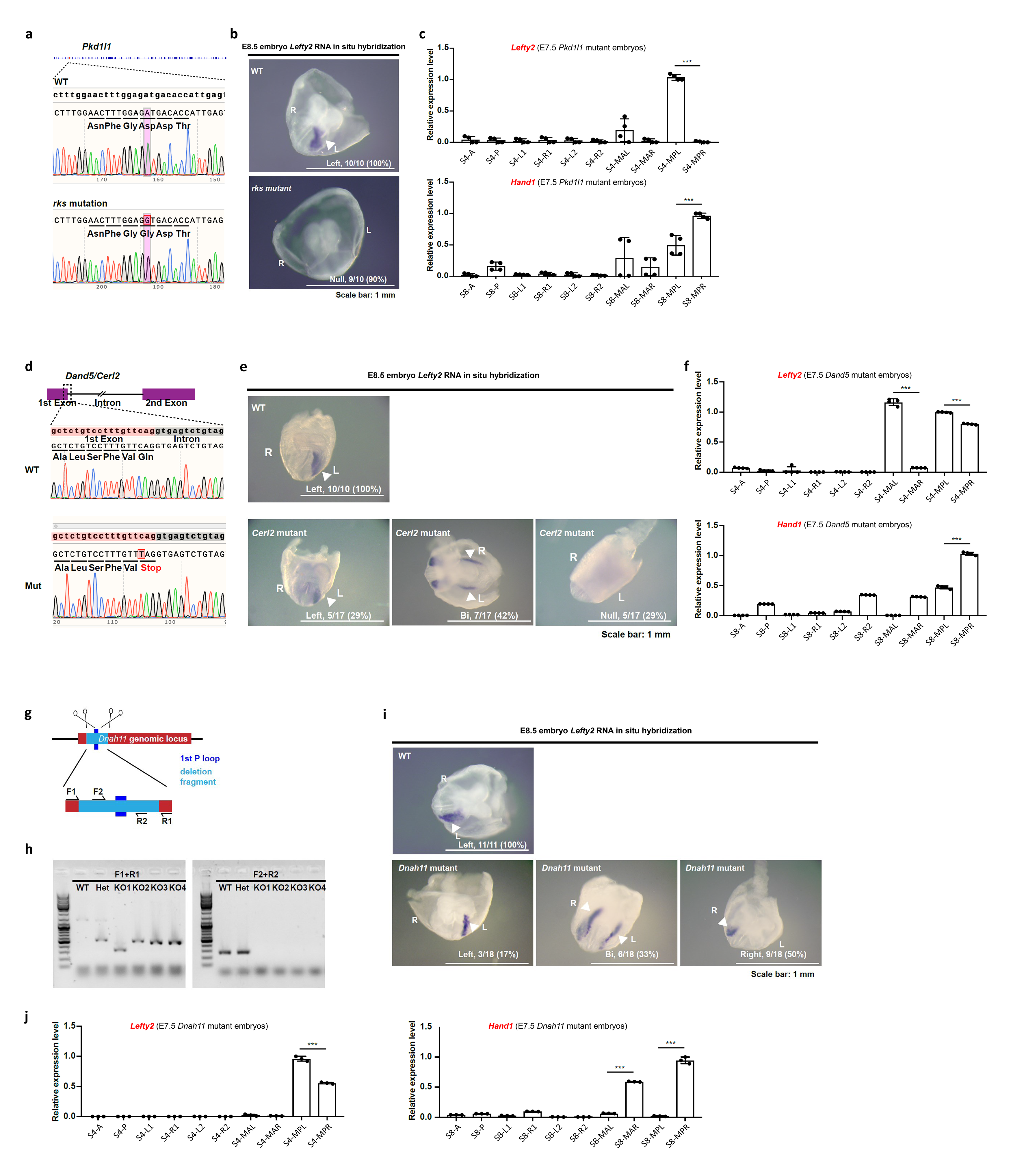
Asymmetric expression of *Lefty2* and *Hand1* persisted in *Pkd1l1*, *Cerl2* and *Dnah11* mutant embryos. **a-c**. *Pkd1l1* mutant embryo: (**a**) Sequencing data of *Pkd1l1* mutant allele. (**b**) Whole-mount in situ hybridization of *Lefty2* in E8.5 mutant embryo. WT, wild type. L, left; R, right. Expression pattern: Null, no expression. (**c**) Real-time PCR showing the relative level of *Lefty2* and *Hand1* expression in E7.5 mutant embryos. **d-f**. *Cerl2* (*Dand5*) mutant embryo: (**d**) Sequencing data of *Cerl2* mutant allele. (**e**) Whole-mount in situ hybridization of *Lefty2* in E8.5 mutant embryo. WT, wild type. L, left; R, right. Expression pattern: left; Bi, bilateral; Null, no expression. (**f**) Real-time PCR showing the relative level of *Lefty2* and *Hand1* expression in E7.5 mutant embryos. **g-j**. *Dnah11* (*iv*) mutant embryo: (**g**) Schematic of the gene editing strategy. (**h**) Genotyping results of *Dnah11* mutant embryo. (**i**) Whole-mount in situ hybridization of *Lefty2* in E8.5 mutant embryo. WT, wild type. L, left; R, right. Expression pattern: left; Bi, bilateral; Null, no expression. (**j**) Real-time PCR showing the relative level of *Lefty2* and *Hand1* expression in E7.5 mutant embryos.

**Extended Data Fig. 17.**
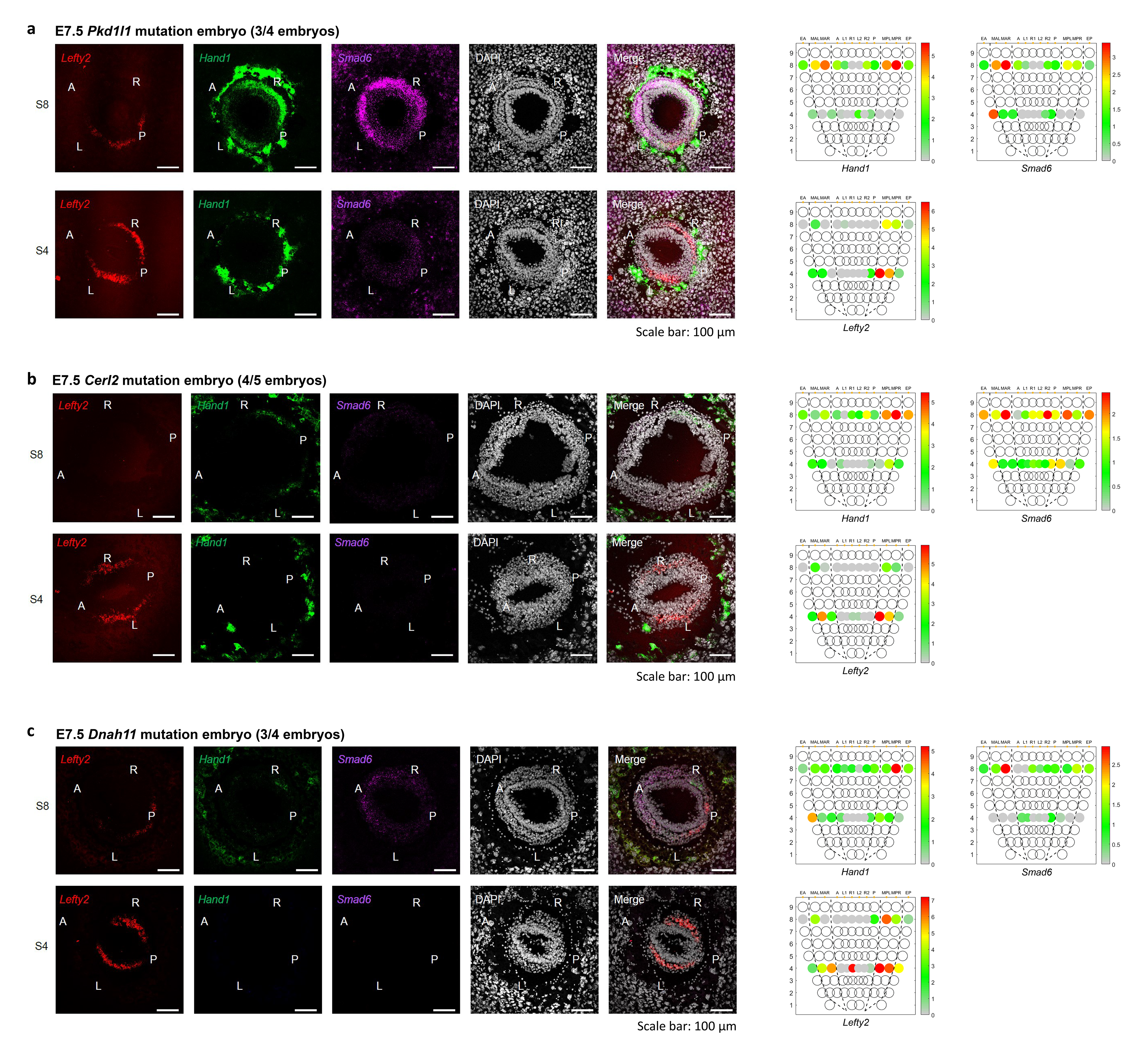
Left-right asymmetry persists in E7.5 mutant embryos. **a-c**. Asymmetric pattern of expression of *Lefty2*, *Hand1* and *Smad6* in E7.5 *Pkd1l1* (**a**), *Cerl2* (**b**) and *Dnah11* (**c**) mutant embryos revealed by RNAscope analysis (immunofluorescence images, A, anterior; P, posterior; L, left; R, right) and Geo-seq analysis (corn plots).

## Notes

### Competing Interest Statement

The authors have declared no competing interest.

